# The lnc-FANCI-2 intrinsically restricts RAS signaling in HPV16-infected cervical cancer

**DOI:** 10.1101/2024.02.01.578320

**Authors:** Haibin Liu, Lulu Yu, Vladimir Majerciak, Thomas Meyer, Ming Yi, Peter F. Johnson, Maggie Cam, Douglas R. Lowy, Zhi-Ming Zheng

**Author notes:** Co-first authors (Haibin Liu, Lulu Yu). Center for Emerging Infectious Diseases, Wuhan Institute of Virology, and Center for Biosafety Mega-Science, Chinese Academy of Sciences, Wuhan, Hubei, 430207, China (Haibin Liu).

## Abstract

We recently discovered the expression of a long noncoding RNA, lnc-FANCI-2, coinciding with cervical lesion progression from CIN1, CIN2-3 to cervical cancer. Viral E7 of high-risk HPVs and host transcription factor YY1 are two major factors promoting lnc-FANCI-2 expression. To explore possible roles of lnc-FANCI-2 in HPV-induced cervical carcinogenesis, we ablated the expression of *lnc-FANCI-2* in the HPV16-positive cervical cancer cell line, CaSki. Knock-out (KO) single cell clones expressed HPV16 oncogenes normally but displayed altered cell morphology and proliferation when compared with the parental cells. Proteomic profiling of cytosolic and secreted proteins from the parental and KO cells showed that lnc-FANCI-2 regulates expression of a subset of cell soluble receptors responsible for cell signaling, epithelial mesenchymal transition, and IFN responses. RNA-seq analyses revealed that, relative to the parental cells, lnc-FANCI-2 KO cells exhibited significantly increased RAS signaling but decreased IFN pathways. In the KO cells, phosphorylated Akt and Erk1/2, two important RAS pathway effectors, were increased more than 3-fold, accompanied by increase of IGFBP3, MCAM, VIM, and CCND2 (cyclin D2) and decrease of RAC3 expression. High levels of lnc-FANCI-2 and lower levels of MCAM in cervical cancer patients are associated with improved survival. We found that lnc-FANCI-2 interacts specifically with 32 host proteins, including H13, HNRH1, K1H1, MAP4K4, and RNPS1, and knockdown of MAP4K4 in CaSki cells led to an increase of phosphorylation of Erk1/2 and somewhat Akt. In summary, a key function of lnc-FANCI-2 is to intrinsically regulate RAS signaling to impact cervical lesion progression and affect cervical cancer prognosis.

**Significance:** Expression of lnc-FANCI-2 is related to cervical lesion progression. Knock-out (KO) or knock-down (KD) of lnc-FANCI-2 expression in HPV16-positive cervical cancer cells CaSki significantly increases RAS signaling, phosphorylation of Akt and Erk1/2, and the epithelial mesenchymal transition factors. lnc-FANCI-2 KO also regulates the expression of a subset of cell soluble receptor proteins IGFBP3 and MCAM. A high level of lnc-FANCI-2 and lower level of MCAM in cervical cancer patients are associated with improved survival. lnc-FANCI-2 in CaSki cells interacts specifically with 32 host proteins, including MAP4K4. KD of MAP4K4 expression in CaSki cells led to an increase in phosphorylation of Erk1/2 and Akt. Thus, one lnc-FANCI-2 function is to intrinsically regulate RAS signaling to impact cervical lesion progression.

## INTRODUCTION

Cervical cancer is a leading cause of mortality in women worldwide. It is the second most diagnosed cancer among women, with an estimated 661,000 new cases worldwide each year, and the third most frequent cause of cancer-related death, accounting for 348,000 deaths annually [1]. Persistent infections in the cervical epithelium with the cancer-associated alpha human papillomaviruses (high-risk HPVs), in particular HPV16 and HPV18 with the sustained expression of viral oncoproteins, E6 and E7, contribute to squamous epithelial cell tumorigenesis [2–4]. Approximate 31.3% of high-grade neoplastic cervical lesions progress to invasive cervical cancer in 30 years [5], while most lesions resolve spontaneously. The progression to cervical cancer typically takes 15 to 20 years after initial infection (https://www.who.int/news-room/fact-sheets/detail/cervical-cancer). These observations suggest that active host defense systems play important roles in preventing malignant transformation of neoplastic cervical lesions, and additional factors besides E6 and E7 are necessary for cervical cancer development. Indeed, one tumor suppressor network, the Fanconi anemia (FA) pathway [6], can be activated by HPV-induced DNA damage. FA proteins including FANCA, FANCD2 and FANCI are elevated upon HPV infection [7–9] and the activated FA pathway restricts HPV replication and guards host genome stability [10]. FANCD2 mutation stimulates HPV16 and HPV31 genome amplification and also promotes cervical and vaginal cancer development in HPV16 E7 transgenic mouse [11, 12].

HPV-positive cervical cancer tissues in general exhibit wild-type p53, wild-type K-RAS, and no overexpression of RAS genes [13–16]. Viral oncoprotein E7 plays a major role in the immortalization of primary epithelial cells mainly by inactivation of pRb family of proteins [17], whereas E6 mediates degradation of tumor suppressor p53 and activation of human telomerase reverse transcriptase (hTERT) transcription [18–20]. However, high-risk E6 and E7 are necessary but not sufficient to transform fully the immortalized cells into malignant cells [21–25] and additional oncogenic stress is needed. For example, the HRAS^G12V^, a constitutively activated RAS GTPase mutation, triggers malignant transformation of E6/E7-expressing keratinocytes [26–29]. Normal RAS regulates cell proliferation, cell differentiation and cell adhesion through cellular signal transduction from growth-factor receptors such as insulin-like growth factors (IGFs) and insulin-like growth factor binding proteins (IGFBP1-6) [30–32]. Thus, overactive RAS signaling might facilitate transformation of E6/E7-expressing keratinocytes. In transgenic mice, HPV16 E6 and E7 can promote the development of spontaneous epithelial skin tumors, but not spontaneous tumors of the reproductive tract [33–36]. Prolonged estrogen treatment is required [37]. Estrogen receptor (ER) activation stimulates the mitogen-activated protein kinase (MAPK/Erk) and phosphoinositide 3-kinase (Pl3K/Akt) pathways, both of which are mediated by RAS [38]. However, the molecular mechanism underlying estrogen-induced cervical cancer in E6 and E7 transgene mice is not fully understood.

Long non-coding RNAs (lncRNAs) are RNAs over 200 nucleotides (nt) in length lacking a coding capacity for translation into functional proteins. Although several lncRNAs have been proposed to function in diverse biological processes, evidence is still lacking to support the functionality of the majority of lncRNAs [39, 40].

We recently reported an increased level of lnc-FANCI-2 expression in high-risk HPV-positive cervical intraepithelial neoplasia (CIN) and invasive cervical cancer (ICC) tissues [8]. At least 14 RNA isoforms of lnc-FANCI-2 are expressed by usage of two alternative promoters, alternative RNA splicing, and alternative selection of two polyadenylation sites from the genomic locus adjacent to the FANCI on Chr5. Ironically, the lnc-FANCI-2 was wrongly annotated by NCBI as a *MIR9-3HG* in the recently updated RefSeq database (https://www.ncbi.nlm.nih.gov/refseq/) and two alternative lnc-FANCI-2 promoters identified for expression of lnc-FANCI-2 are ∼10 kb downstream of the non-expressible miR9-3 gene [8]. Both lnc-FANCI-2 and FANCI are up-regulated simultaneously in neoplastic cervical lesions and cervical cancer by high-risk HPV infections. E6 but, in particular E7 are responsible for the enhanced expression of lnc-FANCI-2, the transcription of which is also regulated by YY1[8]. HPV infection increases YY1 levels but decreases the expression of p53-dependent miR-29a which targets the YY1 3ʹ UTR. Viral E7 interacts with YY1 and facilitates YY1 transactivation of lnc-FANCI-2 promoter [8]. In situ hybridization has revealed that lnc-FANCI-2 is preferentially cytoplasmic, but its function in HPV infected cells has not been characterized. In this report, we demonstrate that lnc-FANCI-2 in HPV16-infected cells controls RAS signaling by interaction with MAP4K4 and other RNA-binding proteins. Ablation of lnc-FANCI-2 in the cells promotes RAS signaling and phosphorylation of Akt and Erk. High levels of lnc-FANCI-2 and low level of MCAM expression in cervical cancer patients correlate with improved survival, indicating that lnc-FANCI-2 plays a critical role in regulating RAS signaling to affect cervical cancer progression and patient outcomes.

## RESULTS

### Differential expression of lnc-FANCI-2 in cervical cancer tissues and its derived cell lines

As lnc-FANCI-2 expression is up-regulated along with cervical lesion progression by high-risk HPV infections [8], we further verified the increased expression of lnc-FANCI-2 in cervical cancer tissues by RNAscope single molecule RNA *in situ* hybridization (RNA-ISH) using a lnc-FANCI-2 antisense probe spanning over the major isoform of lnc-FANCI-2 nt 359-1713 region (GenBank MT669800.1). We found both cytoplasmic and nuclear locations of the increased lnc-FANCI-2 ISH signals within the tumor nest in HPV16-infected cervical squamous cell carcinoma tissues, but not much so in its adjacent tissue areas (Fig. 1A/B). The increased expression of lnc-FANCI-2 became obvious in human foreskin keratinocytes with HPV16 or HPV18 infection when compared to the HFK cells without HPV infection (Fig. 1C).

**Fig. 1.**
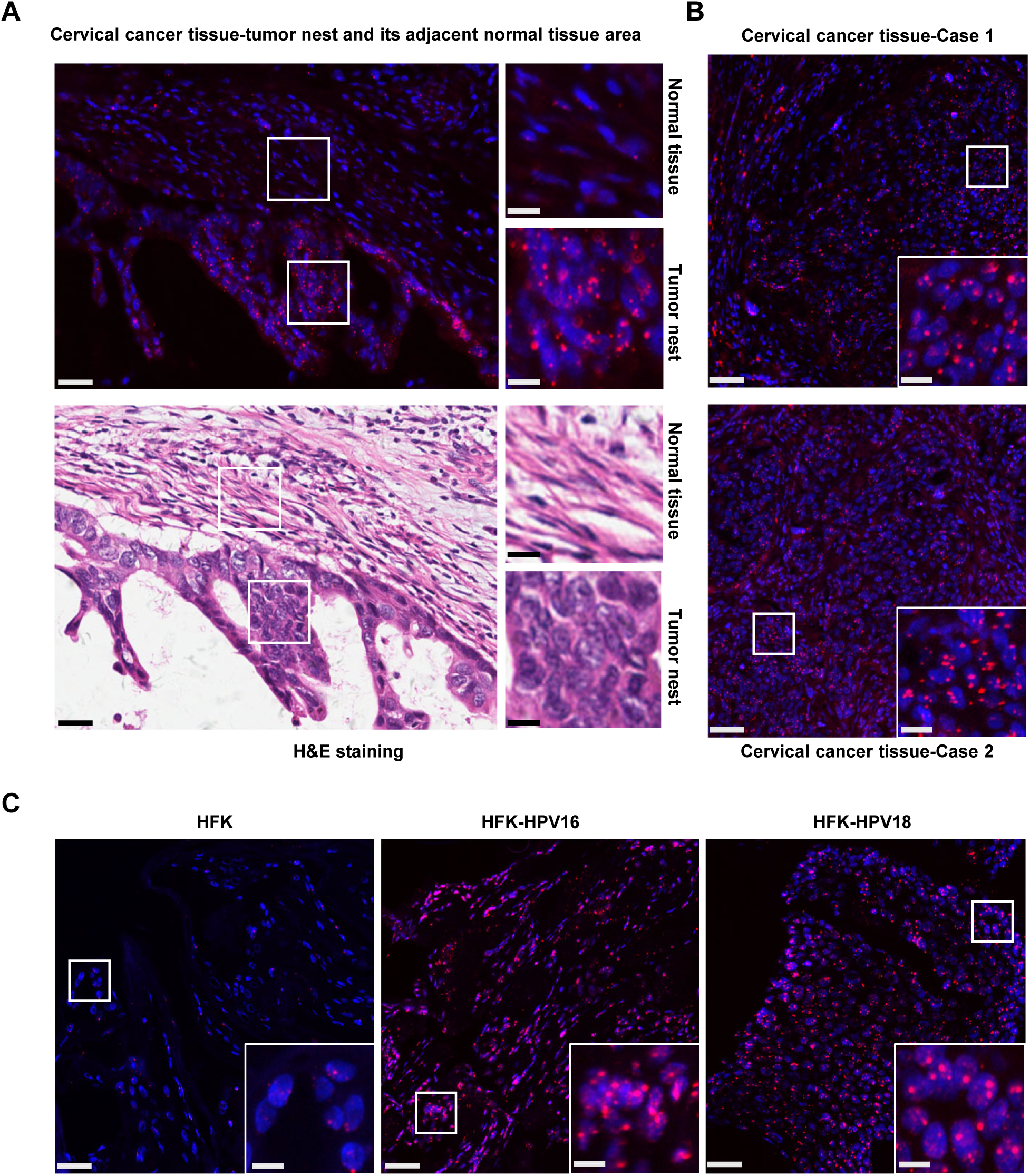
Increased expression of lnc-FANCI-2 in HPV16-infected cervical squamous cell carcinoma tissues and HPV16- and HPV18-infected raft culture tissues. (A and B) Expression lnc-FANCI-2 (red) in HPV16^+^ cervical squamous cell carcinoma tissues were examined by RNAscope RNA-ISH analysis. Nuclei were stained with DAPI (blue). The corresponding regions (A) in H&E staining are shown for the adjacent region from the tumor nest and histotype. (C) Human foreskin keratinocyte (HFK)-derived raft cultures without (HFK) or with HPV16 or HPV18 infections (HFK-HPV16 or HFK-HPV18) were examined at day 10 for lnc-FANCI-2 RNA by RNAscope RNA-ISH analysis. Scale bars: 25 μm in the original figures and 10 μm in the zoomed insets.

However, we subsequently demonstrated the high expression of lnc-FANCI-2 in two HPV16-infected cervical cancer cell lines (SiHa and CaSki) and two HPV16-infected low-grade cervical lesion-derived cervical keratinocyte subclone lines (W12 20861 and W12 20863 cells), but not in two HPV18-infected cervical cancer cell lines (HeLa and C4II cells), nor other HPV-negative cell lines of HCT116 (colorectal cancer cells), BCBL-1 (body cavity B lymphoma cells), HEK293 (Ad5 E1/E2-immortalized human kidney cells), and HaCaT (spontaneously immortalized human epidermal cells) (Fig. S1A). Interestingly, the high expression of lnc-FANCI-2 was observed in a HPV-negative cervical cancer cell line C33A cells with mutations of both p53 and RB genes [41] (Fig. S1A). Northern blot confirmed the increased expression of lnc-FANCI-2 in C33A cells but no expression in HeLa cells (Fig. S1B).

Subcellular distribution of lnc-FANCI-2 was characterized by cell fractionation and Western blot assays using nuclear SRSF3 and cytoplasmic GAPDH protein as an indication of fractionation efficiency (Fig. S1C). By RT-qPCR analysis of the fractionated total RNA, we demonstrated that lnc-FANCI-2 RNA is mainly cytoplasmic in HPV16-positive CaSki cells, but nuclear in HPV16-positive SiHa and HPV-negative C33A cells (Fig. S1C). The differential subcellular distributions of lnc-FANCI-2 RNA from CaSki to SiHa and C33A cells (Fig. S1D) were further confirmed by RNAscope RNA-ISH using the lnc-FANCI-2 antisense probe described above.

### lnc-FANCI-2 regulates proliferation of HPV-transformed cervical cancer cells

lnc-FANCI-2 RNA is transcribed mainly from a proximal promoter TSS2, but also from an alternative minor, distal promoter TSS1. Two highly conserved YY1 binding motifs upstream of the TSS2 are essential for the TSS2 transcriptional activity [8] (Fig. 2A). To elucidate the function of lnc-FANCI-2 in high-risk HPV-infected cervical cancer cells, we knocked out (KO) lnc-FANCI-2 expression in HPV16-positive CaSki cells using CRISPR/Cas9 deletion of either a 3-kb promoter region encompassing both TSS1 and TSS2 or an 86-bp region containing two YY1-binding motifs in the TSS2 promoter (Fig. 2A). Two tested gRNAs with high KO efficiency were selected and cloned into a modified CRISPR/Cas9 expression vector to express both 5ʹ-specific gRNA and 3’-specific gRNA simultaneously for efficient genome editing [42]. CaSki cells stably transfected with the dual gRNA expression vector were generated. Through a serial dilution, several single cell clones were isolated and verified for homozygous deletion by PCR screening. PCR genotyping indicated successful homozygous deletion of the promoter or YY1-binding motifs, respectively (Fig. 2B).

**Fig. 2.**
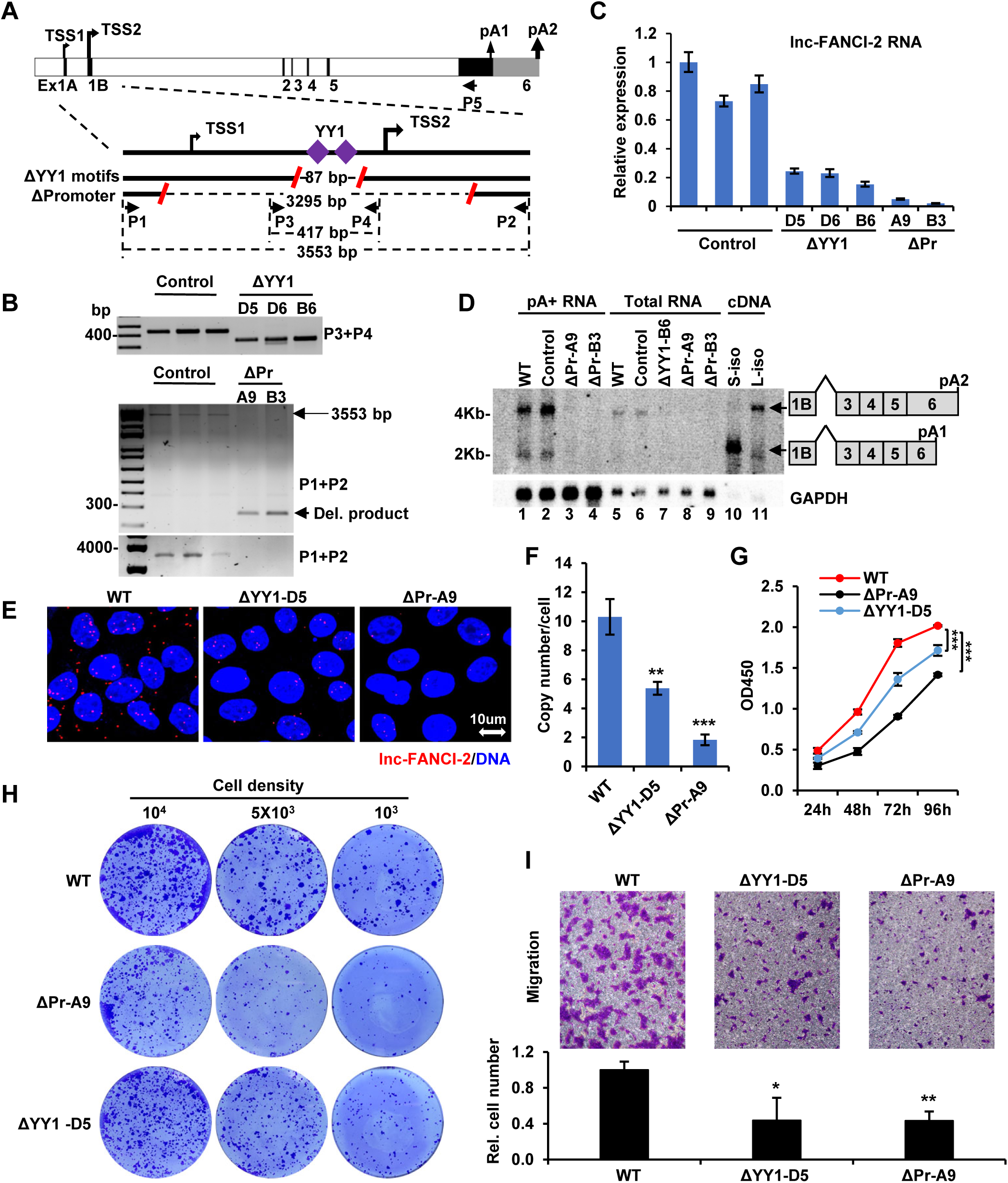
Knockout (KO) of lnc-FANCI-2 in CaSki cells affects cell proliferation, colony formation and migration. (A) Diagram and KO strategies of lnc-FANCI-2 gene. On the figure top is the lnc-FANCI-2 gene structure and its alternative transcription start sites (TSS) and polyadenylation sites (pA). TSS2 or pA2 with a heavier arrow are predominately used for lnc-FANCI-2 expression. The lower figure part shows KO strategies. Red slashes represent gRNA-targeted sites to create genomic DNA deletions by CRISPR-Cas9 technology. Deletion of two YY1-binding motifs (ΔYY1) led to a deletion of 86-bp DNA fragment. Deletion of lnc-FANCI-2 promoter (ΔPr) led to delete a ∼3.3-kb promoter region. Primers (P1-P4) used for PCR screening were shown as black arrows. (B) PCR screening of single cell clones with homozygous lnc-FANCI-2 KO, with indicated primer sets diagramed (A). Single cell clones selected from the cells transfected with an empty vector served as control (Ctrl) cells. (C) Evaluation of lnc-FANCI-2 KO efficiency in the selected single cell clones (B) by RT-qPCR. (D) Northern blot validation of lnc-FANCI-2 KO efficiency from the individual single cell clones or the parental (WT) CaSki cells on polyA^+^ RNA (enriched from 100 μg of total cell RNA, lanes 1-4) or 10 μg of total cell RNA (lanes 5-9). Total cell RNA (2 μg) of HEK293T cells with ectopic expression of two isoforms (short, S or long, L) of lnc-FANCI-2 cDNA served as a control. Antisense oligo probe P5 (A) labeled with ^32^P was used for hybridization. GAPDH RNA served as a loading control and hybridized with a GAPDH-specific oligo probe. (E-F) RNA in situ hybridization (RNA-ISH) validation of lnc-FANCI-2 KO efficiency by RNAscope technology. Two single cell clones were examined with red color for lnc-FANCI-2 and blue color for the nucleus (E). Bar graphs show the copy number of lnc-FANCI-2 per cell in the WT, ΔYY1-D5 or ΔPr-A9 CaSki cells (each averaged from 200 cells) (F). (G-I) Cell proliferation, colony formation and migration of WT, ΔYY1-D5 or ΔPr-A9 cells show requirements of lnc-FANCI-2 expression for cell proliferation, colony formation, and cell invasion of CaSki cells. Representative pictures are shown from three separate experiments of cell proliferation by CCK8 assay (G), cell colony formation under 1% methylcellulose solution (H), and cell migration by transwell assay (I) of WT, ΔYY1-D5 or ΔPr-A9 cells. Cell colonies were stained with crystal violet as described in the procedure. Numbers of the invaded cells crossed the transwell membrane were quantified and shown as bar graphs (I). *, P< 0.05; **, P<0.01; ***, P<0.001 by two tailed Student *t* test.

Loss of lnc-FANCI-2 expression in the single-cell clones was examined by RT-qPCR (Fig. 2C). When compared to all control clones transfected and selected from an empty vector containing no gRNA, all single cell clones with deletion of YY1-binding motifs (ΔYY1) showed 70-80% reduction of lnc-FANCI-2 expression, whereas the single cell clones with deletion of both TSS1 and TSS2 promoters (ΔPr) showed >90% reduction (Fig. 2C). Northern blot analysis confirmed the decrease in levels of the two major lnc-FANCI-2 isoforms (Fig. 2D), with an abundant 4-kb lnc-FANCI-2 RNA derived from a distal polyadenylation site pA2 and a less abundant 2-kb of lnc-FANCI-2 RNA derived from a proximal polyadenylation site pA1[8] (Fig. 2A and 2D). Reduced lnc-FANCI-2 RNA expression was also confirmed by RNAscope RNA-ISH (Fig. 2E and 2F). The parental (WT) CaSki cells expressed ∼10 copies of lnc-FANCI-2 RNA per cell [8], whereas the copy number of lnc-FANCI-2 RNA in the ΔYY1-D5 cells dropped to ∼5 copies and to ∼2 copies per cell in the ΔPr-A9 cells (each averaged from 200 cells).

Although the WT CaSki cells preferentially grow as distinct cluster cell islands, the lnc-FANCI-2 KO cells displayed a dispersed cell growth pattern and some cells exhibited an irregular or spindle-like cell morphology (Fig. S2). The KO cells also grew slower than the parental CaSki cells (Fig. 1G), formed fewer colonies with reduced size (Fig. 1H), and showed decreased migration capacities (Fig. 1I), with ΔPr-A9 cells having a more severe phenotype than ΔYY1-D5 cells. Subsequently, the ΔPr-A9 and ΔPr-B3 cells were further examined for their HPV16 E6 and E7 expression and the ΔPr-A9 for cell senescence. We found both ΔPr-A9 and ΔPr-B3 cells, when compared with parental WT CaSki cells, exhibited some small, variable changes in the expression of E6, E7, p53 (E6 downstream target) and E2F1 (E7 downstream target) proteins (Fig. S3A), most likely from sampling in the assays, but the ΔPr-A9 cells displayed significant increase in cell senescence (Fig. S3B). Altogether, the data indicate that lnc-FANCI-2 RNA is important for cell growth, colony formation, and cell migration.

### lnc-FANCI-2 regulates the expression and secretion of cell soluble receptors

Considering that the altered expression of cell membrane proteins and secreted factors in the lnc-FANCI-2 KO cells might contribute to the observed different cell morphology and growth properties, we performed a Proteome Profiler Human sReceptor Array analysis to examine possible changes in expression of 105 well-characterized soluble protein receptors being involved in cell signaling using total cell lysates and cell culture supernatants from ΔPr-A9 and WT CaSki cells.

Using 30% cut-off (FC −/+0.3), we identified from the ΔPr-A9 cell lysate nine proteins with increased expression compared to the WT CaSki cells, including PODXL2, ECM1, NECTIN2, MCAM, ADAM9, ADAM10, CDH5, ITGA5 and NOTCH1, and six proteins with decreased expression including ITGB6, CDH13, LGALS3BP, TIMP2, ADAM8 and SCARF2 (Fig. 3A and 3B). We also found from the ΔPr-A9 cell culture supernatant five proteins with increased expression, including ADAM9, NECTIN2, ADAM10, CDH5 and ECM1, and the decreased expression of four proteins, including CRELD2, SDC1, SDC4 and TIMP2 (Fig. 3A and 3B). By immunoblot assays, we verified selectively the increase in PODXL2, MCAM, and ECM1 and the decrease in ADAM8 and TIMP2 from the ΔPr-A9 cell lysates and the increase in ECM1 and the decrease in TIMP2 in the ΔPr-A9 cell culture supernatant (Fig. 3C). As ADAM8 is proteolytically processed into two protein isoforms for cell adhesion [43] and migration[44], we confirmed by immunoblot the decreased expression of all three sizes of ADAM8 protein in the ΔPr-A9 cell lysate over the WT CaSki cells (Fig. 3C).

**Fig. 3.**
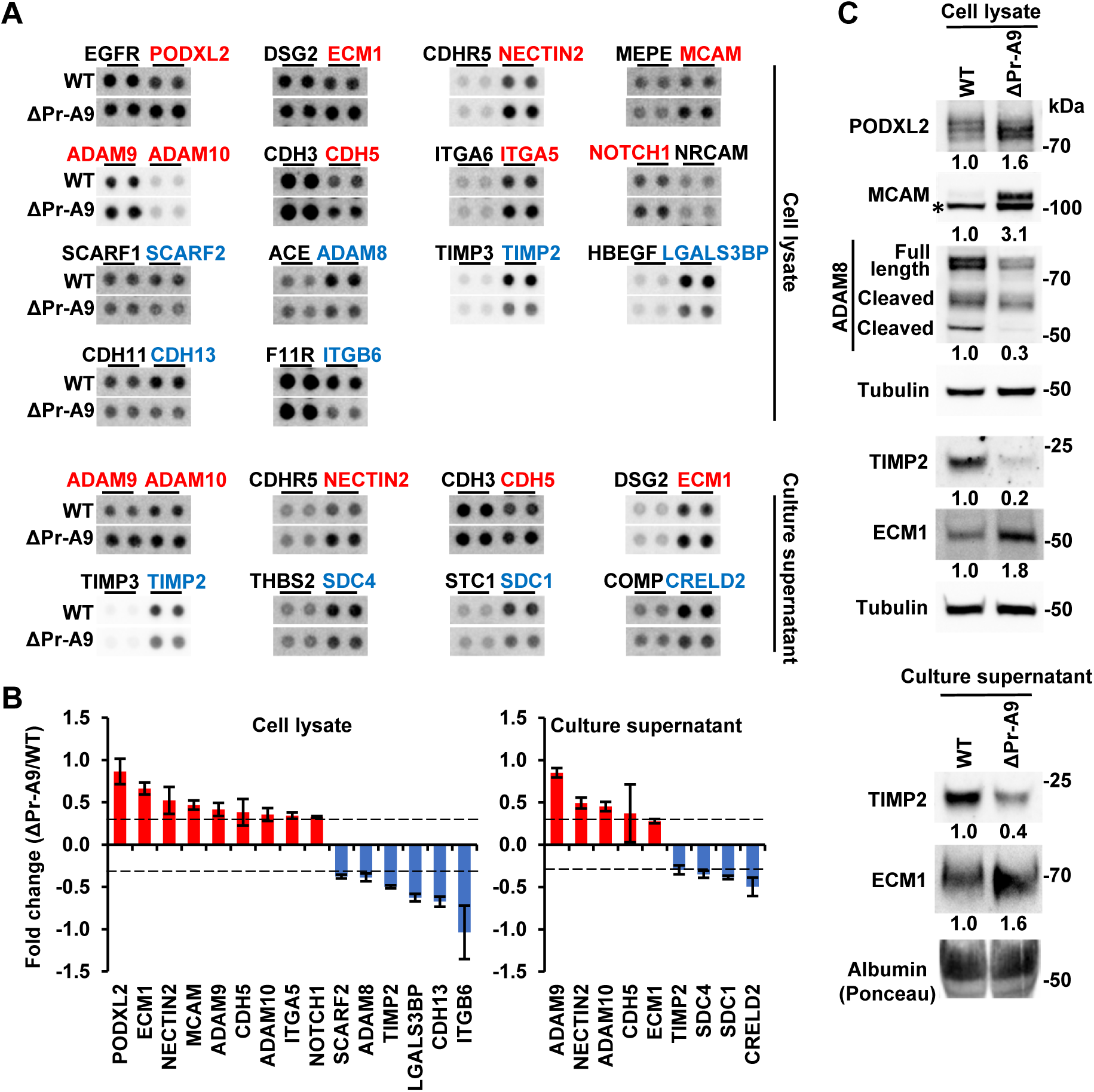
KO of lnc-FANCI-2 in CaSki cells affects expression of cellular soluble receptors. (A) Dot blots show differentially expressed soluble receptors in ΔPr-A9 cells when compared to those in the WT CaSki cells. A Proteome Profiler Human sReceptor Array was used to examine 105 cellular soluble receptors. Label in red, upregulated receptors; label in blue, downregulated receptors. (B) Quantification of differentially expressed and/or released soluble receptors (A) in ΔPr-A9 cells when compared to those in the WT CaSki cells. Error bar represents standard deviation of replicates in each proteome array. The dash line indicates the threshold of fold change (FC) with a score above or below the threshold (-/+ 0.3 FC) being determined as differentially expressed and/or released receptors. (C) Validation of several differentially expressed soluble receptors by immunoblot analysis using specific antibodies. Tubulin served as an internal loading control. * non-specific band in MCAM immunoblot.

### lnc-FANCI-2 regulates the expression of genes involved in RAS signaling

We next conducted genome-wide RNA-seq analyses of WT and lnc-FANCI-2 KO cells (four samples/group) to determine the transcriptomic consequences of lnc-FANCI-2 deficiency. We obtained ∼120 million mappable RNA reads to the human reference genome hg38 from each sample. Analysis of RNA-seq reads-coverage map by Integrative Genomics Viewer (IGV) showed ∼90% reduction of lnc-FANCI-2 expression in ΔPr-A9 cells and ∼72% decrease in ΔYY1-D5 cells compared to the WT CaSki cells (Fig. 4A), consistent with the results shown in Fig. 2C-F.

**Fig. 4.**
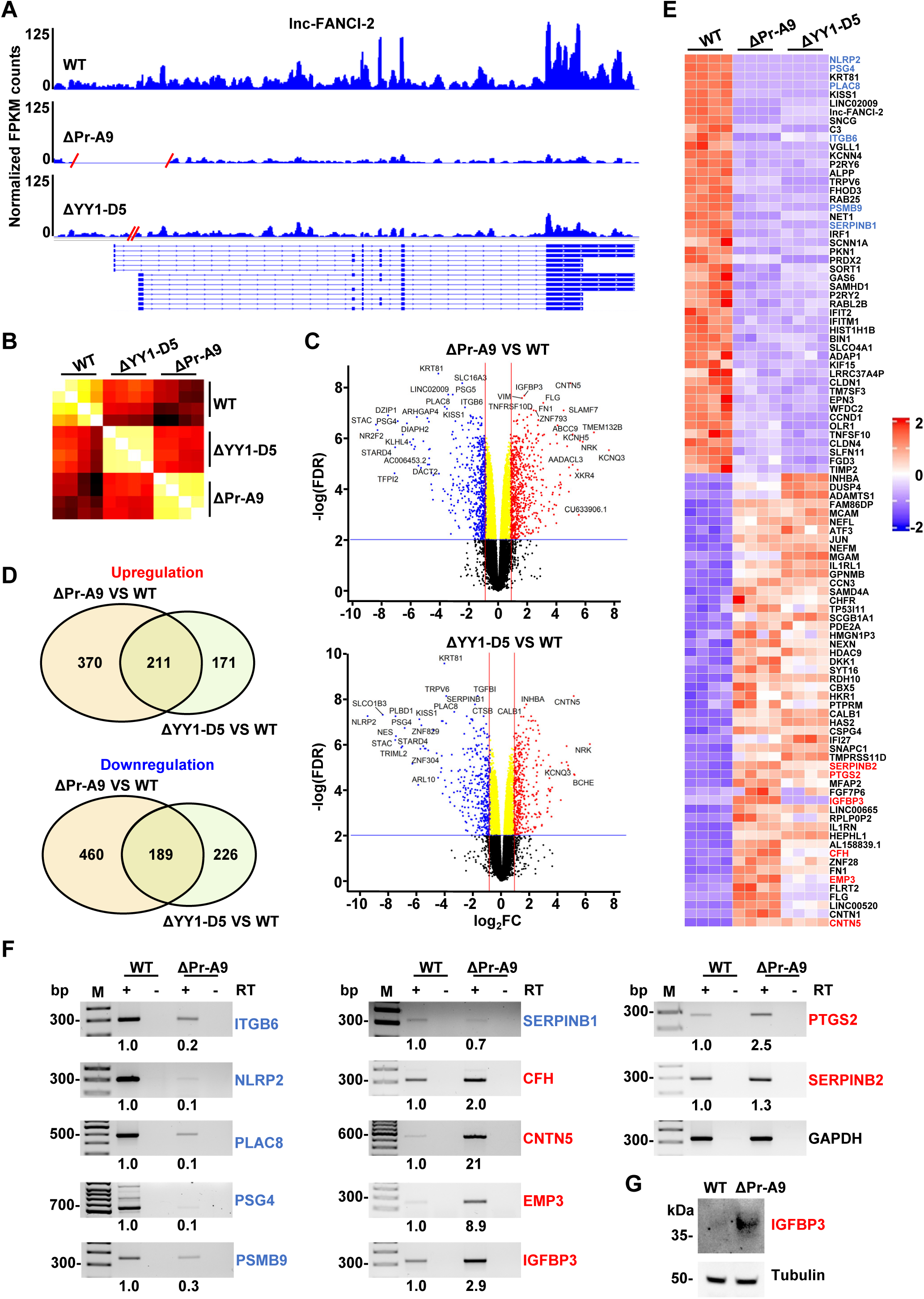
Transcriptomic effect of lnc-FANCI-2 KO in CaSki cells by RNA-seq analysis. (A) RNA-seq reads-coverage maps by IGV showing the expression levels of lnc-FANCI-2 from two KO cell clones, ΔPr-A9 and ΔYY1-D5, to the WT CaSki cells. one representative coverage profile of four in each type of cells is shown by IGV. FPKM, fragments per kilobase of transcript per million. Red slashes represent the deleted genomic region, with validated fourteen lnc-FANCI-2 RNA isoforms shown below [8]. (B) Similarity in heatmap comparison among ΔPr-A9, ΔYY1-D5 and WT cells was generated using Limma-normalized counts from each sample by Pearson complete linkage method. (C) Volcano plot visualization of differentially expressed genes (DEGs) in ΔPr-A9 and ΔYY1-D5. The genes with the most significant change in the increased (red) or decreased (blue) expression are indicated. FC, fold change. (D) Venn diagram depicting the overlapped DEGs between ΔPr-A9 and ΔYY1-D5 cells over the WT CaSki cells. (E) Heatmap shows overlapped DEGs with FPKM ≥7 in ΔPr-A9 and ΔYY1-D5 cells when compared with the WT CaSki cells. (F) Validation of selective 12 upregulated or downregulated DEGs shown in the heatmap by RT-PCR with gene-specific primers in the presence (+) or absence (-) of reverse transcriptase (RT). GAPDH served as an internal RNA control. Relative expression level of each gene was calculated based on the band density after normalizing to GAPDH RNA band, with the expression level in the WT CaSki cells setting as 1. M, a 100-bp DNA marker. (G) Validation of the upregulated expression of IGFBP3 in ΔPr-A9 cells over the WT CaSki cells by immunoblot analysis.

Hierarchical clustering analysis showed more global transcriptional similarity between ΔYY1-D5 and ΔPr-A9 cells (Fig. 4B). By applying a threshold of fold change (FC) ≥ 1.8 or FC ≤ −1.8 with FDR ≤ 0.01, we found 1230 genes in ΔPr-A9 and 797 genes in ΔYY1-D5 cells with significantly differential expression relative to the WT CaSki cells expressing ∼15890 genes. Nine (*Italic*) of 15 (60%) soluble receptor proteins in cell lysates (PODXL2, ECM1, *<u>NECTIN2</u>, <u>MCAM</u>, <u>ADAM9</u>*, CDH5, ADAM10, *<u>ITGA5</u>*, *<u>NOTCH1</u>*, SCARF2, ADAM8, *<u>TIMP2</u>*, *<u>LGALS3BP</u>*, *<u>CDH13</u>*, and *<u>ITGB6</u>*) exhibited consistent changes in expression by both RNA-seq and protein array assays. The most significantly affected genes are shown in Volcano plots (Fig. 4C and Table S1). Among these, 211 were upregulated and 189 were downregulated in both ΔPr-A9 and Δ YY1-D5 cells (Fig. 4D and Table S2). By applying more stringent criteria with FPKM ≥7 in at least one of four RNA-seq samples as a cutoff, we profiled the genes with large expression differences in both ΔPr-A9 and ΔYY1-D5 cells. The results displayed in the heatmap of Fig. 4E included 52 upregulated, and 47 downregulated genes excluding lnc-FANCI-2. Subsequently, we selectively verified by RT-PCR the decreased expression of ITGB6, NLRP2, PLAC8, PSG4, PSMB9, and SERPINB1 and the increased expression of CFH, CNTN5, EMP3, IGFBP3, PTGS2, and SERPINB2 in ΔPr-A9 cells (Fig. 4F), as well as the increased expression of IGFBP3 protein, a RAS signaling driver [30–32], in ΔPr-A9 cells by immunoblot (Fig. 4G).

By performing the Gene Set Enrichment Analysis (GSEA) on the Hallmark gene sets, which provide more refined and concise inputs for GSEA [45], we found the most significantly upregulated pathways in both ΔPr-A9 and ΔYY1-D5 cells, when compared with the WT CaSki cells, were KRAS signaling and epithelial mesenchymal transition (EMT). The most significantly downregulated pathways in both ΔPr-A9 and ΔYY1-D5 cells over the WT CaSki cells were interferon gamma (IFN-γ) and interferon alpha (IFN-α) responses (Fig. 5A). GSEA plots for ΔPr-A9 shows 29 of 111 genes, including IGFBP3 [30–32], in RAS signaling and 39 of 130 genes in EMT (Fig. 5B and 5C, Table S3) were significantly upregulated, whereas 37 of 139 genes in IFN-γ response and 24 of 71 genes in IFN-α response were significantly downregulated (Fig. 5D and 5E, Table S3). Similar enriched gene sets with increased RAS signaling and EMT and decreased IFN-γ and IFN-α responses were observed in ΔYY1-D5 cells (Fig. S4, Table S3).

**Fig. 5.**
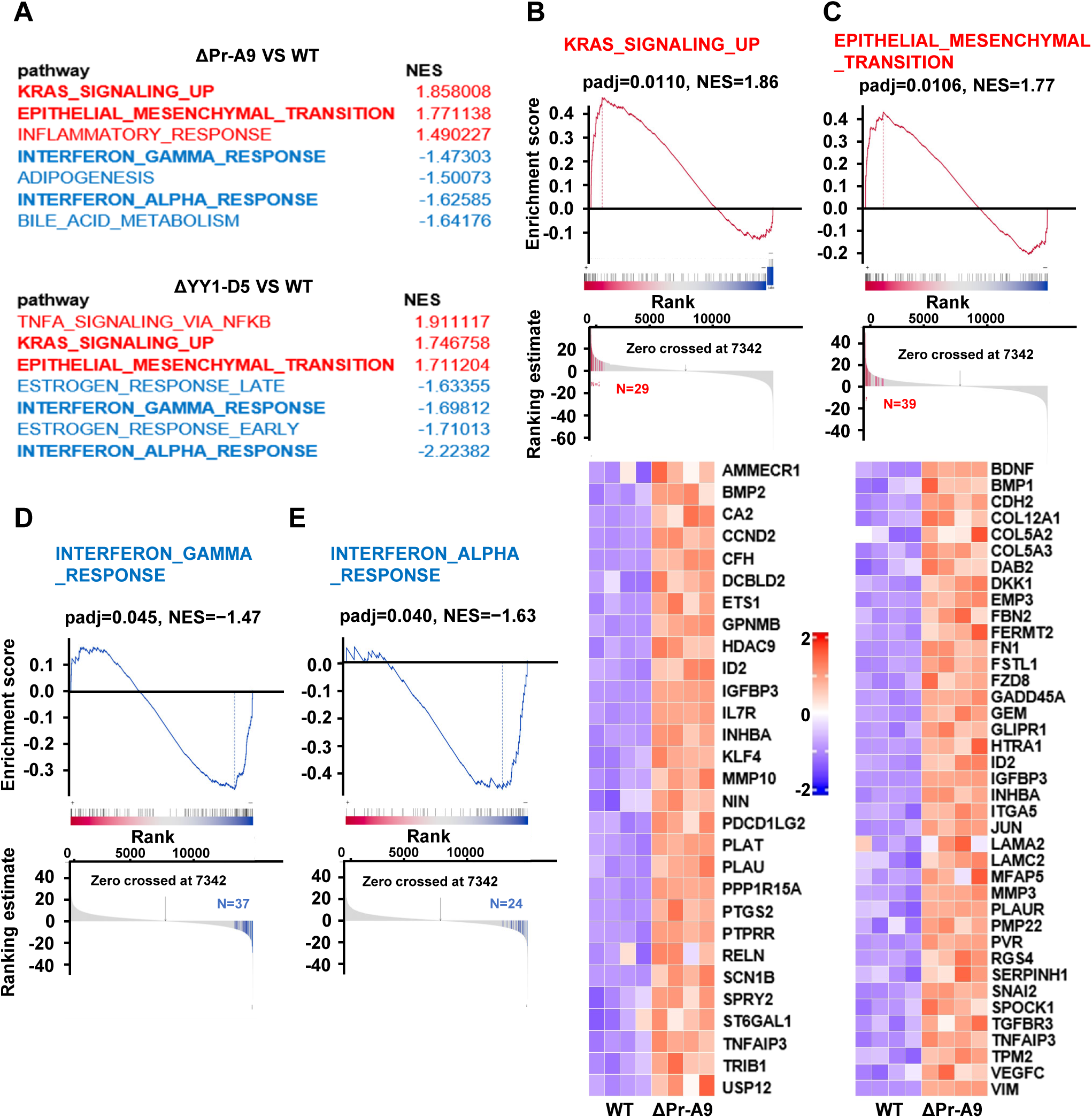
Pathway analyses of DEGs identified by RNA-seq. (A) Top 3 upregulated and top 4 downregulated pathways in ΔPr-A9 and ΔYY1-D5 cells over the WT CaSki cells. Data were generated by Gene set enrichment analysis (GSEA) performed with Hallmark gene sets. NES stands for normalized enrichment score. (B-E) Each GSEA Enrichment plot shows the enrichment score and gene hits enriched in ΔPr-A9 cells using Hallmark gene sets, KRAS_SIGNALING_UP (B), EPITHELIAL_MESENCHYMAL_TRANSITION (C), INTERFERON_GAMMA_ RESPONSE (D) or INTERFERON_ALPHA_RESPONSE (E). padj, adjusted p value; Zero crossed 7342, the middle number of total genes in the GSEA at ranking. The heatmaps below the enrichment plots (B and C) visualize the genes enriched in respective pathways of KRAS_SIGNALING_UP (B) and EPITHELIAL_MESENCHYMAL_TRANSITION (C) in ΔPr-A9 cells when compared to the WT CaSki cells.

Since ΔPr-A9 cells exhibit higher KO efficiency of lnc-FANCI-2 and a higher number of differentially expressed genes (DEGs) and show more severe phenotypic changes in cell proliferation, migration, and colony formation than ΔYY1-D5 cells, the ΔPr-A9 cells were primarily used for further focused studies on RAS signaling in this report.

### lnc-FANCI-2 regulates RAS GTPase activities to phosphorylate RAS signaling effectors Akt and Erk

RAS activation triggers two major downstream signal transduction pathways, Raf/Mek/Erk and PI3K/Akt (Fig. S5), to transduce signals from extracellular stimuli to the cell nucleus where specific effector genes are activated for their corresponding functions [30–32, 46, 47]. To further confirm the GSEA data and verify that two major signaling pathways of RAS are activated, we first examined and compared the RAS GTPase activities in ΔPr-A9 cells and the WT CaSki cells. We found a significant increase in RAS GTPase activity in ΔPr-A9 cells (Fig. 6A, top bar graphs), indicating that the endogenous lnc-FANCI-2 RNA in CaSki cells suppresses not only the expression of IGFBP3 (Fig. 4F and 4G), the most abundant IGFBP, but also RAS activation (Fig. 6A, top bar graphs). Since ΔPr-A9 cells had undergone long term selection and adaption during single cell screening, the increased RAS GTPase activity might result from selection pressure for cell survival. However, by transient siRNA knockdown (KD) of lnc-FANCI-2 expression in the WT CaSki cells, we obtained a similar, albeit weaker, increase of RAS GTPase activity (Fig. 6A, lower bar graphs). This result suggests that the increased RAS GTPase activity in ΔPr-A9 cells was unrelated to the persistent selection pressure, but rather the deficiency of lnc-FANCI-2.

**Fig. 6.**
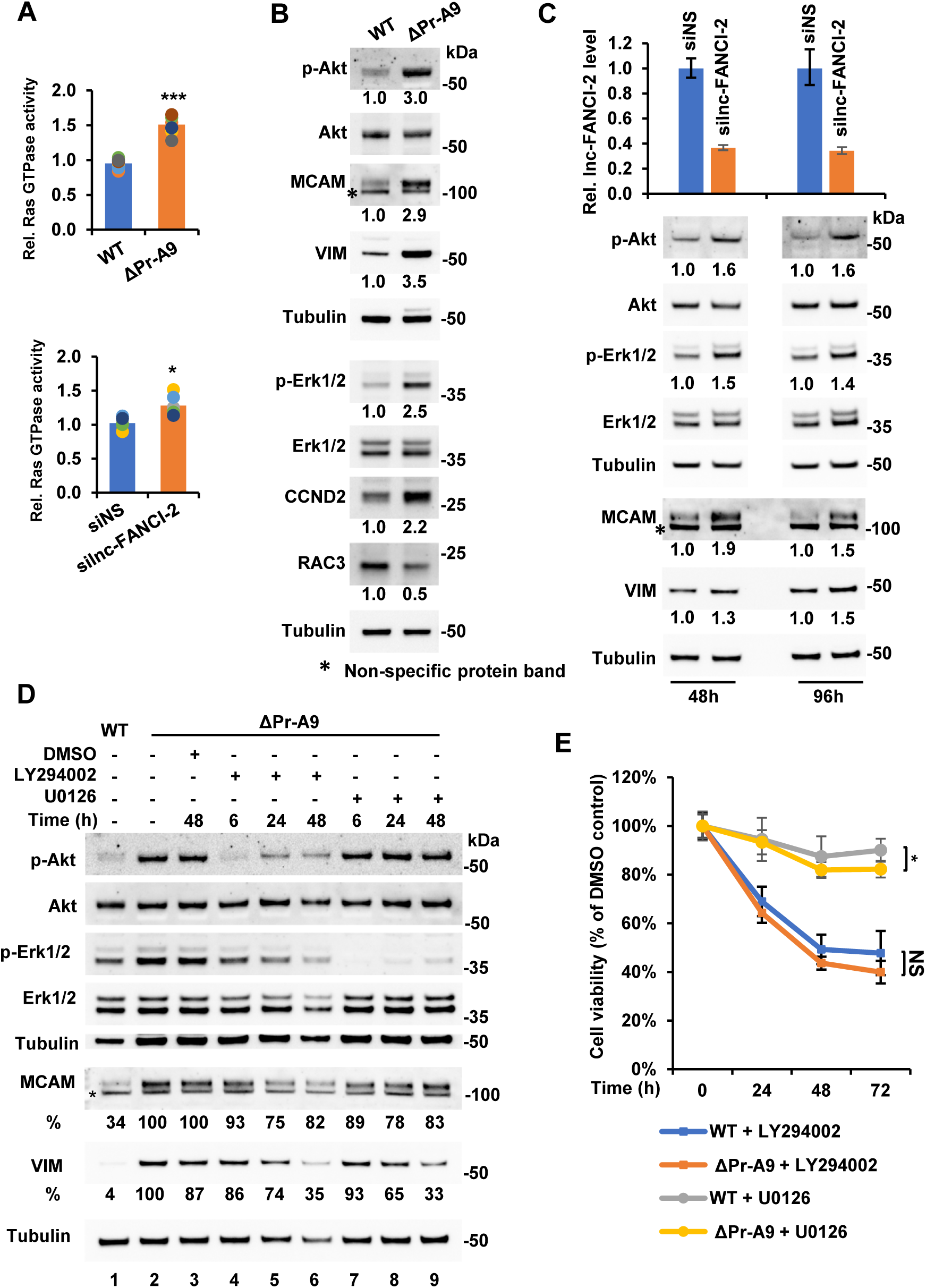
CaSki cells with lnc-FANCI-2 KO exhibit activation of RAS signaling pathway. (A) CaSki cells with lnc-FANCI-2 KO (top, ΔPr-A9 cells) or knockdown (KD, lower) by lnc-FANCI-2-specific siRNAs display increased RAS GTPase activity when compared to the WT CaSki cells (top) or the WT cells treated with a nonspecific control siRNA (siNS). The relative RAS GTPase activity was measured by a RAS GTPase Chemi ELISA assay. *, P< 0.05; ***, P<0.001 by two tailed Student *t* test. (B and C) Selective validation of increased expression of RAS signaling-related downstream genes in lnc-FANCI-2 KO ΔPr-A9 cells (B) over the WT CaSki cells or lnc-FANCI-2 KD CaSki cells (C) over the WT cells treated with a nonspecific control siRNA (siNS). Relative expression of the indicated genes in ΔPr-A9 cells in comparison to the WT cells (B) or in the WT CaSki cells treated for 48 h and 96 h with a nonspecific control siRNA (siNS) or a siRNA specifically targeting the lnc-FANCI-2 exon 3 (C) were immunoblotted using the corresponding antibodies as indicated. Tubulin served as an internal control for each blot. The level of p-Akt or p-Erk was calculated by normalizing to total Akt or Erk protein and the other proteins by normalizing to tubulin. The protein level in the WT cells or WT cells treated with siNS set as 1. The KD efficiency of lnc-FANCI-2 RNA (C, top bar graphs) was examined by RT-qPCR. (D) Effect of blocking RAS signaling on the expression of MCAM and VIM using PI3K inhibitor LY294002 (20 μM) and MEK inhibitor U0126 (10 μM). The protein at each time point was immunoblotted with a corresponding antibody. The level of MCAM or VIM was calculated after normalizing to tubulin. The protein level in ΔPr-A9 cells without the inhibitor was set as 100%. *, nonspecific protein band. (E) The time dependent cell viability of the WT CaSki and ΔPr-A9 cells in the presence of 20 μM LY294002 or 10 μM U0126. Data was obtained in each time point after normalizing to the cells treated with DMSO. The mean ± SD at each data point was calculated from 6 samples combined from two independent experiments.

Next, we examined activation of the PI3K/Akt and Raf/Mek/Erk pathways [48–50]. We observed a 3-fold increase in phosphorylated Akt (p-Akt) and 2.5-fold increase of p-Erk1/2 (p44/p42 MAPK), after being normalized to total Akt and Erk levels, in ΔPr-A9 cells over the WT CaSki cells (Fig. 6B). These results were confirmed in ΔYY1-D5 cells (Fig. S6A). The increased p-Akt and p-Erk (mostly p-Erk2/p42) was accompanied by elevated expression of MCAM, VIM, and CCND2, and decreased expression of RAC3 (Fig. 6B and Fig. S6A). These randomly chosen potential RAS effector proteins facilitate soluble receptor functions, cell proliferation, and EMT. Moreover, transient siRNA knockdown of lnc-FANCI-2 in the WT CaSki cells also led to the increased levels of p-Akt and p-Erk (Fig. 6C) and increased expression of MCAM and VIM at 48 h and 96 h post-transfection (Fig. 6C). These observations indicate that a transient loss of lnc-FANCI-2 in CaSki cells is sufficient to trigger RAS signaling. The same siRNA transfection in HeLa cells, a lnc-FANCI-2 negative cell line (Fig. S1A and S1B), exhibited no effect on p-Akt or p-Erk1/2 levels (Fig. S6B), but in SiHa cells containing mainly nuclear lnc-FANCI-2 (Fig. S1C and S1D) enhanced the expression of p-Akt and p-Erk1/2 (Fig. S6B), verifying the specific effect of lnc-FANCI-2 depletion on RAS signaling pathways independently of predominant cellular distribution of lnc-FANCI-2. The lnc-FANCI-2 KO-mediated increase in p-Akt and p-Erk was sensitive to PI3K inhibitor LY294002 [51] (Fig. 6D, lanes 4-6) and MEK1/2 inhibitor U0126 [52] (Fig. 6D, lanes 7-9), respectively. The inhibitors also decreased the expression of VIM and MCAM and cell proliferation, in particular by MEK1/2 inhibitor U0126 (Fig. 6D and 6E). The effects of LY294002 on cell proliferation were similar from ΔPr-A9 to the WT CaSki cells (Fig. 6E).

### lnc-FANCI-2 inhibits the expression of IGFBP3 and MCAM (CD146 or MUC18)

Given that IGFBP-3 is the most abundant IGFBP among all six IGFBP members in potentiation of IGF action and PI3K/AKT activities [30] and a reduced expression of IGFBP-3 mRNA level is associated with progression to cervical cancer [53], we further investigated the effect of lnc-FANCI-2 on the expression of IGFBP3 as lnc-FANCI-2 effector gene. As shown in Fig. 4F-G and Fig. S7A, the increased RNA and protein expression of IGFBP3 appears in the lnc-FANCI-2 KO ΔPr-A9 cells over the parental WT CaSki cells, indicating a suppressive effect of lnc-FANCI-2 on IGFBP3 expression. This suppressive function of lnc-FANCI-2 was further confirmed in lnc-FANCI-2 rescue experiments in the ΔPr-A9 cells (Fig. 7A and 7B). By transient expression of one major isoform of lnc-FANCI-2 RNA (a-PA2, GenBank: MT669800.1) [8] in the ΔPr-A9 cells, lnc-FANCI-2 (red) and IGFBP3 RNA (green) at 24 h post transfection were detected by RNAscope RNA-ISH using each specific antisense RNA probe (Fig. 7A). We demonstrated that the cells with rescued expression of cytoplasmic lnc-FANCI-2 RNA displayed much reduced expression of cytoplasmic IGFBP3 RNA (Fig. 7B).

**Fig. 7.**
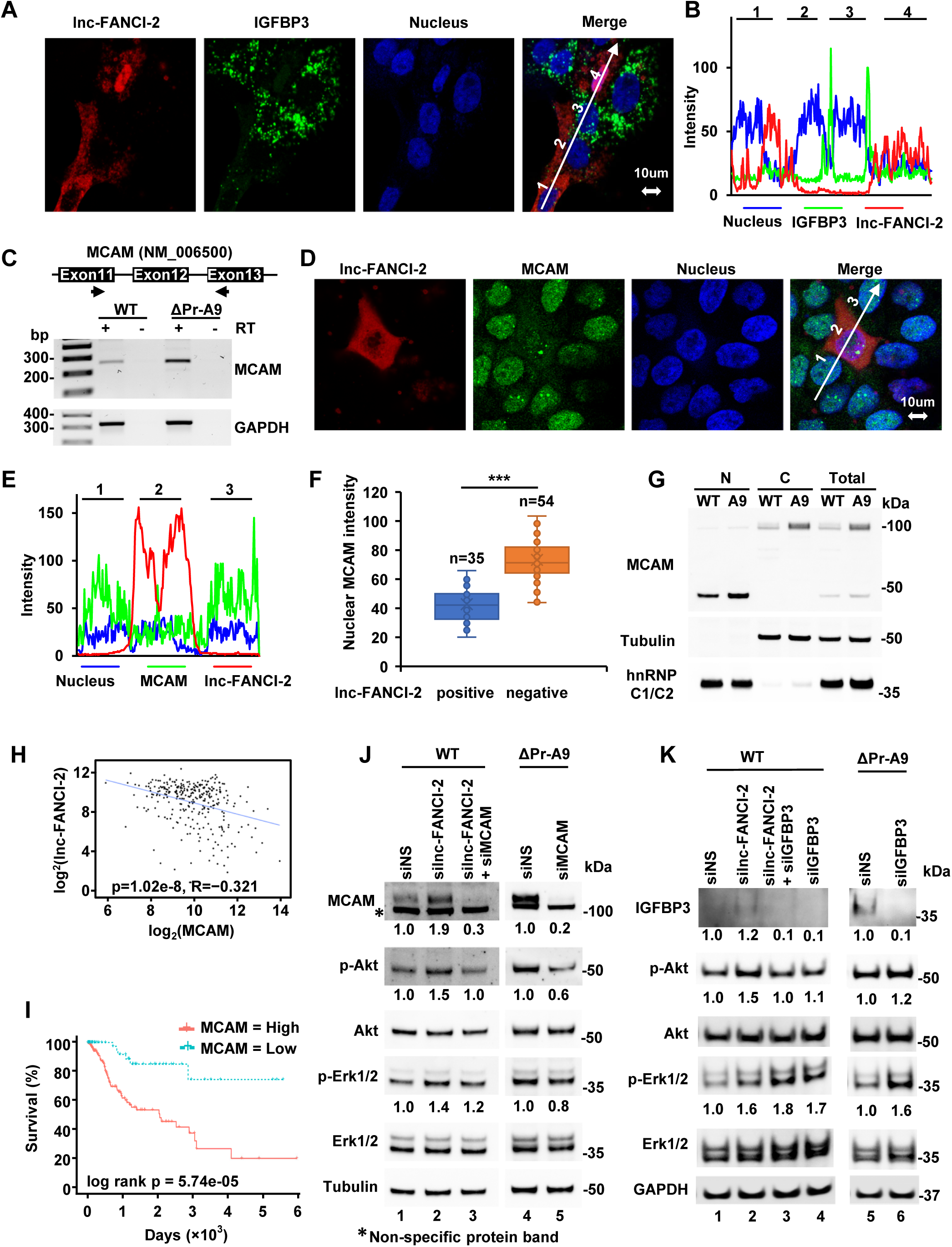
lnc-FANCI-2 is suppressive to the expression of IGFBP3 and MCAM. (A and B) Transient rescue expression of lnc-FANCI-2 in Δpr-A9 cells inhibits the expression of IGFBP3. ΔPr-A9 cells were transfected with a major isoform lnc-FANCI-2a-PA2 (GenBank ACC. No. MT669800.1) cDNA plasmid. lnc-FANCI-2 in red and IGFBP3 RNA in green were detected by RNAscope RNA-ISH at 24h post transfection using each specific antisense RNA probe and imaged by confocal microscopy (A). Expression levels of lnc-FANCI-2 RNA and IGFBP3 RNA in four neighboring cells (B) were measured by signal intensity of a line crossing over the stained cells in A (white line arrow). (C) Validation of differential expression of MCAM RNA in ΔPr-A9 cells by RT-PCR in the presence (+) or absence (-) of reverse transcriptase (RT). One pair of primers with one primer at the exons 11 and the other at exon 13 of MCAM RNA (NM_006500) were used. GAPDH served as a loading control. (D) The major isoform lnc-FANCI-2a-PA2 repressed the expression of MCAM. ΔPr-A9 cells were transfected with lnc-FANCI-2a-PA2 cDNA plasmid. lnc-FANCI-2 RNA in red was detected by RNAscope ISH and MCAM protein in green was detected by IF with an anti-MCAM antibody. (E) Expression levels of lnc-FANCI-2 RNA and MCAM protein in three neighboring cells were measured by signal intensity of a line crossing over the stained cells (D, white line arrow). (F) Calculation of the expression levels of MCAM in lnc-FANCI-2 positive cells (n=35) and lnc-FANCI-2 negative cells (n=54) by fluorescent intensity. (G) Subcellular MCAM distributions in the nucleus (N) and cytoplasm (C) of WT CaSki cells and ΔPr-A9 by immunoblot analysis. Fractionation efficiency and sample loading were controlled by cytoplasmic (represented by tubulin) and nuclear (represented by hnRNP C1/C2) proteins. (H-I) Correlation and survival analysis of lnc-FANCI-2 and MCAM expression with cervical squamous cell carcinoma (CESC) cases from the cancer genome atlas (TCGA) datasets by GEPIA web server (http://gepia.cancer-pku.cn/). The negative correlation at R= −0.321 (with p-value=1.02e-08) of Inc-FANCI-2 with MCAM in cervical cancer patients was obtained from the RNA-seq data from the TCGA CESC tumors downloaded from the TCGA data portal (https://portal.gdc.cancer.gov/). Only primary solid tumor samples (n=304 after exclusion of 2 metastatic samples and 3 normal samples) were subjected to analysis, with the data showing as a Scatter plot (H). Kaplan-Meier plot with log rank test p-value at 5.74e-05 (I) shows MCAM as a biomarker for poor prognosis of cervical cancer survival with a lower quartile group cutoff. RNA-seq and survival data are derived from the TCGA CESC cancer patients. (J/K) AKT and ERK phosphorylation is partially regulated by MCAM and IGFBP3 in CaSki cells. KD of MCAM (J) and IGFBP3 (K) expression in WT parental CaSki or ΔPr-A9 cells was treated with MCAM siRNA or IGFBP3 siRNA along with or without lnc-FANCI-2 siRNA for 48 h. Expression levels of individual proteins were immunoblotted using the corresponding antibodies. The level of MCAM, p-Akt, or p-Erk1/2 was calculated after normalizing to tubulin. The protein level in a non-targeting siRNA (siNS) control set as 1.

As shown in Fig. 4E and Fig. S7B, the increased RNA expression of MCAM (CD146 or MUC18) [54] appears to be transcriptional or posttranscriptional in both ΔPr-A9 and ΔYY1-D5 cells. We confirmed by RT-PCR the lnc-FANCI-2 KO-mediated increase of MCAM RNA expression in ΔPr-A9 cells (Fig. 7C). A rescue experiment was further performed by transient expression of one major isoform of lnc-FANCI-2 RNA (a-PA2, GenBank: MT669800.1) [8] in ΔPr-A9 cells to determine both nuclear and cytoplasmic MCAM expression [55, 56] in individual lnc-FANCI-2 expressing cells. ΔPr-A9 cells ectopically expressing diffused cytoplasmic lnc-FANCI-2 RNA, as detected by RNAscope RNA-ISH, exhibited a marked reduction in nuclear MCAM protein (Fig. 7D-7E). Quantitative analyses of the ΔPr-A9 cells with or without transient lnc-FANCI-2 RNA expression showed significant reduction of nuclear MCAM protein in the 35 cells with rescued lnc-FANCI-2 expression when compared to the 54 cells expressing no lnc-FANCI-2 (Fig. 7F). Due to RNAscope procedures, the membrane-bound and cytoplasmic MCAM protein was mostly removed by protease III treatment when RNA-ISH was performed and thus only the nuclear signal of MCAM protein remained was detectable by IF staining. Using cell fractionation, we detected a cleaved MCAM of ∼46 kDa mainly in the nuclear fraction by Western blot (Fig. 7G). The increased nuclear MCAM expression in ΔPr-A9 cells was proportional to the cytoplasmic and total MCAM levels when compared with the WT CaSki cells (Fig. 7G). The nuclear MCAM signal appeared as a proteolytic cleavage product presumably by the increased metalloprotease ADAM9 and ADAM10 proteins and decreased metalloprotease inhibitor TIMP2 (Fig. 3), as observed in other studies [56].

The significance of an inverse correlation of lnc-FANCI-2 and MCAM in CaSki cells was further investigated by analyzing their expression in 304 cervical cancer samples from the TCGA dataset. There was a significant negative correlation in lnc-FANCI-2 (being wrongly assigned as LINC00925 by NCBI) [8] and MCAM RNA levels (Fig. 7H). The opposite effects of lnc-FANCI-2 and MCAM on cervical cancer survival (Fig. 7I and Fig. S8) show that the cervical cancer patients with a higher level of lnc-FANCI-2 [8] but a lower level of MCAM in the cervical tissue (Fig. S8 and Fig. 7I) exhibited a better survival prognosis.

### Regulatory roles of lnc-FANCI-2-mediated increase of MCAM and IGFBP3 on RAS signaling

To further dissect the lnc-FANCI-2-associated expression of MCAM and IGFBP3 on RAS signaling, we examined MCAM and IGFBP3 on phosphorylation of Akt and Erk1/Erk2 in CaSki cells in the presence or absence of lnc-FANCI-2. As shown in Figure 7J, we found, although KD or KO of lnc-FANCI-2 promotes phosphorylation of Akt and Erk (Fig. 7J, lane 2), that KD of MCAM expression in the parental WT CaSki cells in the absence of lnc-FANCI-2 (Fig. 7J, lane 3) or in ΔPr-A9 cells (Fig. 7J, lane 5) led to reduction of Akt and Erk phosphorylation. These data suggest that MCAM is a lnc-FANCI-2 effector but also could be a trigger of signal transduction as reported [54, 57].

IGFBP3 protein has been viewed as a RAS signaling regulator [30–32], but recent studies show that IGFBP3 has a variety of intracellular ligands involved in many unexpected functions [58, 59]. Thus, KD or KO of lnc-FANCI-2-mediated IGFBP3 expression on RAS signaling in the parental WT CaSki cells or ΔPr-A9 cells was examined by Western blot after siRNA KD of IGFBP3 expression. As shown in Figure 7K, KD of IGFBP3 expression was found to increase phosphorylation of Erk1/2 by ∼70% (lane 4) to ∼60% (lane 6), but not much so for Akt phosphorylation (lanes 4 and 6). Instead, KD of IGFBP3 expression could prevent lnc-FANCI-2 KD-enhanced Akt phosphorylation in the parental WT CaSki cells (Fig. 7K, compare lane 2 and lane 3). These data suggest a separate role of IGFBP3 on phosphorylation of Erk1/2 from Akt in the presence or absence of lnc-FANCI-2.

### MAP4K4 association with lnc-FANCI-2 RNA in CaSki cells regulates RAS signaling and phosphorylation of Akt and Erk

We speculated lnc-FANCI-2 restriction on RAS signaling through interactions with cellular proteins. Subsequently, comprehensive identification of lnc-FANCI-2-binding proteins in the parental WT CaSki cells was performed by an IRPCRP protocol (modified from published ChIRP [60]) in combination with mass spectrometry (Fig. 8A). In this protocol, the lnc-FANCI-2-binding proteins in the parental WT CaSki cells were covalently crosslinked to lnc-FANCI-2 RNA via UV irradiation and pulled down by pooled 30 antisense oligos crossing over the entire lnc-FANCI-2. The RNA extracted from the pulldowns were verified to be lnc-FANCI-2 specific by RT-PCR in the absence (-) or presence (+) of reverse transcriptase (RT) (Fig. 8B). The lnc-FANCI-2-binding proteins in the pulldowns were then subjected to LC-MS/MS analyses. We identified 32 specific lnc-FANCI-2-binding proteins, including H13, HNRH1, K1H1, MAP4K4, and RNPS1 (Fig. 8C, Table S4).

**Fig. 8.**
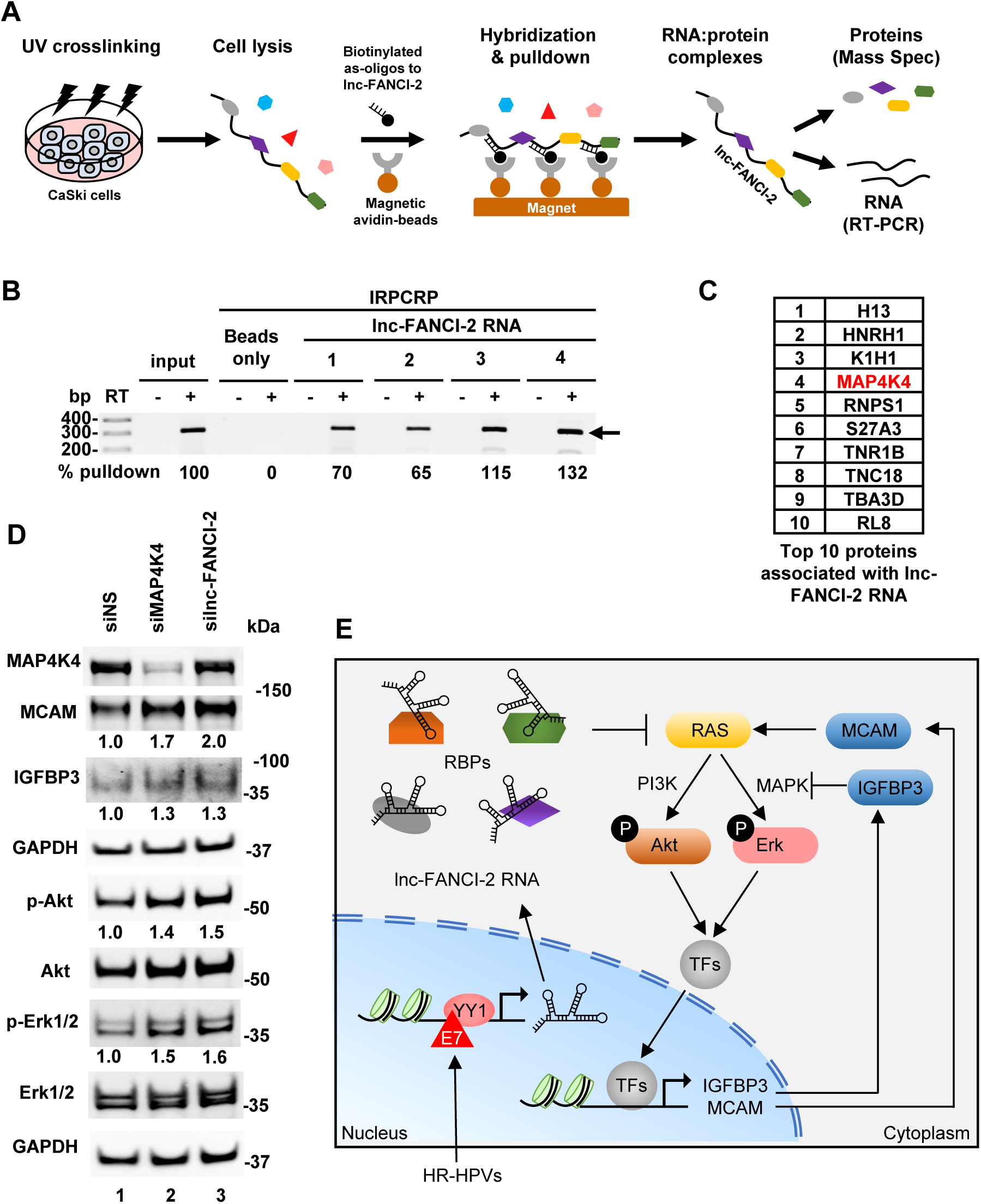
lnc-FANCI-2 interacts with host factors to regulate RAS signaling pathway. (A) The lnc-FANCI-2-associated proteins in the WT CaSki cells were identified by isolation of <u>R</u>NA-<u>p</u>rotein <u>c</u>omplexes using <u>R</u>NA <u>p</u>urification (IRPCRP)-mass spectrometry technology. (B) lnc-FANCI-2 RNA in the IRPCRP-1 and IRPCRP-2 pulldowns had the pooled antisense biotinylated oligos (pool 1 with oligos in even numbers and pool 2 with oligos in odd numbers) immobilized to avidin-beads first before mixed with cell lysates, while the IRPCRP-3 and IRPCRP-4 pulldowns had the oligos pool 1 and 2 separately mixed with cell lysates first before addition to the avidin-beads for the RNA pull-downs. RT-PCR in the absence (-) or presence (+) of reverse transcriptase (RT) was carried out using the RNA isolated from the individual IRPCRP experiments using a primer pair of oHBL5 and oHBL12 (Table S5) specific for lnc-FANCI-2 RNA detection. Beads only (no oligos) IRPCRP experiments served as a negative control. Total RNA from the WT CaSki cells after sonication was used as an input control. The arrow indicates the detected lnc-FANCI-2 RNA. (C) Proteins associated with lnc-FANCI-2 RNA identified from lnc-FANCI-2 IRPCRP pulldowns. A total of 32 proteins were specifically pulled down from lnc-FANCI-2 IRPCRP reactions 1-4 (PSM≥2 from two separate pulldowns, Table S4), with top 10 proteins binding lnc-FANCI-2 shown in the order by the number of identified PSM (peptide spectrum match) in LC-MS/MS. (D) Expression of p-Akt and p-Erk from CaSki WT cells 48 h after siRNA KD of MAP4K4 or lnc-FANCI-2 was immunoblotted by the corresponding antibodies. GAPDH served as a sample loading control. Fold change of the indicated proteins in the cells with KD of MAP4K4 or lnc-FANCI-2 over the cells treated by a non-targeting siRNA (siNS) was calculated after normalizing to GAPDH. (E) A proposed model illustrates how lnc-FANCI-2-protein complexes inhibit the RAS signaling pathway to control Akt/Erk phosphorylation and expression of host genes. RAS signaling can be regulated by integrating external and internal factors. In HR-HPV infected cells, viral oncoprotein E7-YY1 complex transactivates lnc-FANCI-2 expression. By interactions with cellular RBPs, the lnc-FANCI-2-protein complex inhibits RAS signaling. In the absence of lnc-FANCI-2, increased RAS signaling leads to phosphorylation of Akt and Erk and cascaded responses of transcription factors (TFs) and thus regulates the expression of RAS signaling responder genes, such as IGFBP3, MCAM, etc. Consequently, this brings fundamental biochemical and biological processes under control by finetuning the RAS pathway.

MAP4K4, a serine/threonine protein kinase, stimulates cancer cell proliferation, invasion and migration, and is recently characterized as a novel MAPK/ERK pathway regulator [61–63] and negatively regulates RAS signaling by binding to Ras p21 protein activator 1 or RASA1 [64, 65]. Although the exact MAK4K4 binding site on lnc-FANCI-2 was not explored, many enzymes turn out an RNA-binding enzyme [66, 67]. Thus, whether MAP4K4 association with lnc-FANCI-2 could regulate PI3K/Akt or MAPK/Erk signal transduction was investigated in the parental WT CaSki cells. We demonstrated that siRNA KD of MAP4K4 expression in the parental WT CaSki cells, as seen for KD of lnc-FANCI-2, led to ∼40% or more increase of phosphorylation of both Akt and Erk1/2 and increased expression of MCAM and IGFBP3 (Fig. 8D, compare lane 2 to lane 3). Interestingly, siRNA KD of MAP4K4 expression in the lnc-FANCI-2 KO ΔPr-A9 cells had no effect on expression of IGFBP3, only a minimal effect on MCAM, but strongly on pErk1/2 (Fig. S7C). The data suggest that, through RNA-protein interactions, the increased lnc-FANCI-2 RNA in cells and their association with MAP4K4 and other cellular proteins could be one arm to control RAS signaling and gene expression of its effectors (Fig. 8E).

## DISCUSSION

HPV oncoproteins E6 and E7 are necessary but not sufficient for development of cervical cancer. Indeed, early studies indicated that other oncogenic stress, such as activated RAS mutant, is needed to trigger malignant transformation of E6 and E7 expressing cells [21, 26, 68, 69] and tumorigenesis [27, 28, 70–74]. Moreover, E7 has been also reported to repress phosphorylation of Akt and Akt-mediated signaling [75]. In this study, we show that lnc-FANCI-2, whose expression is highly dependent on E7 and YY1 [8], intrinsically restricts RAS GTPase activities and phosphorylation of Akt and Erk presumably by interacting with cellular factors in HPV16-positive CaSki cells. Interestingly, early reports indicated that p-Akt attenuation by E7 could be abolished by introduction of H73E mutation in the E7 CR3 domain [75]. This E7 CR3 domain is also essential to interact with transcription factor YY1 to activate lnc-FANCI-2 transcription [8]. Thus, it would be presumable that the reported p-Akt attenuation by E7 might be mediated by the increased expression of lnc-FANCI-2. More importantly, these findings also suggest that additional oncogenic stress, such as activated RAS mutant, is required to overcome the lnc-FANCI-2 restriction on RAS signaling for the early observed malignant transformation [21, 26, 68, 69] and tumorigenesis [27, 28, 70–74] of E6 and E7 expressing cells. In CaSki cells, loss of lnc-FANCI-2 in this report was found to promote RAS signaling but reduction of IFN responses. However, high RAS-AKT-ERK signaling induces cell senescence and inhibits tumor cell survival as shown in this report and in a lung cancer study [76].

Some lncRNAs function as RAS regulators [77, 78] by their ability to sequester RAS-targeting miRNAs, including MALAT1/ miR-217, RMRP/miR-206, and KRAS1P/miR-143/let-7 [78]. Conceivably, lncRNAs may also regulate RAS signaling through other mechanisms beyond miRNAs. In this report, loss of lnc-FANCI-2 in HPV16-positive CaSki and SiHa cells promotes RAS signaling and expression of RAS signaling effectors (Fig. 8E). By profiling the proteins associated with lnc-FANCI-2 in CaSki cells, we identified a serine/threonine protein kinase MAP4K4 as one of major lnc-FANCI-2-binding proteins that may partially mediate the lnc-FANCI-2 restriction on RAS signaling. MAP4K4 is a negative RAS signaling regulator in the context of the early embryo [62, 64] and lymphatic vascular development by interacting with RASA1 [65]. Indeed, silencing of MAP4K4 in parental WT CaSki cells, like KD or KO of lnc-FANCI-2, led to partially increase of both p-Akt and p-Erk1/2 (Fig. 8D) as reported [64, 65].

Interestingly, KD of MAP4K4 in ΔPr-A9 cells led to further increased expression of p-Erk1/2 but not p-Akt, suggesting a negative correlation of MAP4K4 activity in interaction with lnc-FANCI-2 on Erk1/2 phosphorylation. Thus, our data suggest that MAP4K4 protein may function coordinately with lnc-FANCI-2 RNA in the CaSki cells to restrict one arm but not the other of RAS signaling.

All HR-HPVs are capable of inducing lnc-FANCI-2 expression by experimental infection of keratinocytes [8]. HPV^+^ cervical cancer tissues express high level of lnc-FANCI-2. As expected, all HPV16^+^ cervical cancer and pre-cancer cell lines examined, such as CaSki, SiHa, W12 subclone cells 20861(HPV16^+^, integrated) and 20863 (HPV16^+^, episomal), produce lnc-FANCI-2. Unexpectedly, we found no expression of lnc-FANCI-2 in isogenic HPV18^+^ HeLa and C4II cells, nor in colorectal cancer cell line HCT116, adenovirus 5 DNA-transformed human embryonic kidney cell line HEK293, spontaneously transformed aneuploid immortal keratinocyte cell line HaCaT, and KSHV-infected B cell lymphoma cell line BCBL-1 cells which are all HPV-negative cell lines. It remains to know what negatively regulates lnc-FANCI-2 expression in those cell lines and more HPV18^+^ cell lines should be examined for lnc-FANCI-2 expression. However, we did find that a HPV-negative cervical cancer cell line, C33A cells expressing mutant p53 and mutant pRB [41], produce a high level of lnc-FANCI-2 (Fig. S1A and S1B). Using dual luciferase report assays, we did find lnc-FANCI-2 promoter activity in response to YY1 binding in CaSki and C33A cells, but not in HeLa and HCT116 cells, suggesting the presence of a repressive factor (s) in the latter two cell lines (data not shown). KD of E2F1 had no effect on lnc-FANCI-2 promoter activity in CaSki cells [8]. In the lnc-FANCI-2 producing cells, we noticed that cellular location of lnc-FANCI-2 also varies from cell types, with lnc-FANCI-2 predominantly in the nucleus in SiHa and C33A cells, but in the cytoplasm in CaSki cells, although both cytoplasmic and nuclear lnc-FANCI-2 restrict RAS signaling. This suggests that a variety of nuclear and cytoplasmic functions of lnc-FANCI-2 remains to be explored.

IGFBP3 is a major IGF carrier and enhances both IGF- and EGF-RAS signaling [31, 79] to activate phosphatidylinositol 3-kinase (PI3K)/AKT and the mitogen-activated protein kinase (MAPK)/ERK pathways [80]. Surprisingly, we find a suppressive role of IGFBP3 on Erk phosphorylation, but not much on Akt, in the presence or absence of lnc-FANCI-2 in HPV16-positive CaSki cells. Although ∼99% of circulating IGF-1 is bound to IGFBPs, predominantly to IGFBP-3, the most abundant IGFBP in human serum, case-control studies showed significantly lower IGF-1 and IGFBP-3 serum levels in the patients with invasive cervical cancer over the control group [81, 82]. A reduced expression of IGFBP-3 mRNA level appears to be associated with progression to cervical cancer [53]. Consistently with lnc-FANCI-2’s suppressive effect on IGFBP3 expression in our study, an increased level of lnc-FANCI-2 expression is associated step-wisely with high-risk HPV-positive cervical intraepithelial neoplasia and invasive cervical cancer tissues [8].

It is obvious from our RNA-seq and human soluble receptor array analysis that KO of lnc-FANCI-2 in CaSki cells affects the expression of a large set of genes, including the expression of increased NECTIN2, ADAM9, ADAM10, ITGA5, MCAM, PODXL2, ECM1, but decreased ADAM8, TIM2, and ITGB6, which are responsible for RAS signaling, EMT pathway, and IFN responses. For example, the dysregulation of RAS signaling and ADAM protein activity is implicated in various cancers. ADAM proteins can modulate RAS signaling by cleaving and releasing ligands that activate or inactivate RAS-related pathways [83–86]. Some ADAM proteins are Involved in the migration and invasion of cancer cells, and its loss can promote the degradation of KRAS [87]. ADAMs are essential for extracellular signaling and regulation of cell adhesion and their catalytic activity is strongly correlated to Src-homology 3 (SH3) domain binding of SH3 proteins [85, 86, 88]. In addition to the notified activation of PI3K/Akt and Raf/Mek/Erk pathways, one of the major RAS signaling effectors in lnc-FANCI-2 KO or KD cells was found to be MCAM.

MCAM binds various cellular surface receptors or co-receptors to trigger signal transduction, cell proliferation and motility, tumor angiogenesis, and tumor cell metastasis [54, 57]. It is an integral, highly glycosylated membrane protein with three common protein isoforms [54, 89, 90]. Two membrane-anchored forms with a long (p113) or short (p76) cytoplasmic tail produced by alternative RNA splicing and a proteolytically cleaved soluble form without transmembrane and cytoplasmic regions [54, 90, 91]. In addition to its roles in cell signal transduction [54, 57, 89, 90], the short isoform produced by an intramembrane cleavage may be directed towards the nucleus to regulate gene transcription [56]. We found that CaSki cells express mainly a cytoplasmic form of full-length MCAM (∼113 kDa) and a small nuclear form (∼46 kDa), but not the short ∼76 kDa isoform. Moreover, lnc-FANCI-2 KO cells exhibit high MCAM levels, which was most likely a result of the increased RAS signaling. Consistently, lnc-FANCI-2 and MCAM levels are negatively correlated in cervical cancer patients, and low levels of MCAM and high levels of lnc-FANCI-2 indicate better prognosis. This suggests that increased MCAM might promote cervical tumorigenesis in vivo. Given that the cells with Inc-FANCI-2 KO exhibit increased levels of IGFBP3 and MCAM and the ΔPr-A9 cells with rescued lnc-FANCI-2 RNA expression displayed remarkable reduction of IGFBP3 and MCAM expression, our current model (Fig. 8E) highlights how the HPV16-enhanced expression of lnc-FANCI-2 in cervical cancer cells might function as a negative regulator by binding to various RNA-binding proteins to block RAS signaling and reduce the expression of RAS signaling effectors such as IGFBP3 and MCAM, etc.

In conclusion, this study provides the first evidence that one of lnc-FANCI-2 functions is to maintain epithelial functional integrity by repressing RAS signaling in preventing malignant transformation of neoplastic cervical lesions. In addition, more extensive studies are undergoing to elucidate the role of lnc-FANCI-2 in regulation of IFN responses, which are not covered in this report. Since transactivation of lnc-FANCI-2 expression is largely dependent on host transcription factor YY1 of which expression is regulated by HPV oncoproteins and host miR-29a, dissecting their transcription network on lnc-FANCI-2 and its effectors may further provide novel insights involved in host homeostasis against HPV infection-induced carcinogenesis. Our observations also highlight additional, unexpected players beyond viral E6 and E7 that drive cells toward malignant transformation.

## MATERIALS AND METHODS

### Cell cultures and treatment

CaSki, a HPV16-positive (HPV16^+^) human cervical cancer cell line with wild-type p53, pRb, HRAS, KRAS, and NRAS (https://depmap.org/portal/cell_line/ACH-001336?tab=mutation) was obtained from the American Type Culture Collection (ATCC, Manassas, VA), maintained in Dulbecco’s modified Eagle medium (DMEM) (Thermo Fisher Scientific) supplemented with 10% fetal bovine serum (FBS) and 1% penicillin-streptomycin, and grown in a cell culture incubator at 37°C with 5% CO_2_.

SiHa (HPV16^+^), HeLa (HPV18^+^), C4II (HPV18^+^), C33A (no HPV), and other HPV-negative cell lines, HEK293T (Ad5 E1/E2-immortalized human kidney cells), and HaCaT (spontaneously immortalized human epidermal cells), were all obtained from ATCC and grown in Dulbecco’s modified Eagle’s medium (DMEM) with 10% FBS at 37°C and 5% CO_2_. HCT116 cell line, a colorectal cancer cell line, was a gift from Dr. Bert Vogelstein of Johns Hopkins University and was grown in McCoy’s 5A medium with 10% FBS at 37°C and in 5% CO_2_. W12 subclones 20861 (HPV16^+^, integrated) and 20863 (HPV16^+^, episomal) were a gift from Dr. Paul Lambert of University of Wisconsin, Madison, and grown on mitomycin C-pretreated NIH3T3 feeder cells in F12 medium at 37°C and in 5% CO_2_. BCBL-1 (body cavity B lymphoma cells with KSHV infection) was obtained from the AIDS Research and Reference Reagent Program, Division of AIDS, NIAID, NIH and cultured in RPMI 1640 containing 10% FBS at 37°C and in 5% CO_2_.

CaSki cells were treated with 20 µM phosphoinositide 3-kinase (PI3K) inhibitor, LY294002 (Cell Signaling Technology, Danvers, MA, #9901) or 10 µM mitogen-activated protein kinase kinase (MEK) inhibitor, U0126 (Cell Signaling Technology, Danvers, MA, #9903) and examined for their proliferation and viability by CCK-8 assay and RAS effector expression by Western blot.

### Raft tissues and cervical tissue sections

Raft tissues with or without HPV16 or HPV18 infection and cervical cancer tissue sections from Zhejiang University Women’s hospital were described in our previous reports [8, 92].

### siRNAs, plasmid constructions and transfections

Synthetic double-stranded siRNA targeting the lnc-FANCI-2 transcript is listed in a supplementary Table S5. Human MAP4K4 siRNA (#M-003971), human IGFBP3 siRNA (#M-004777) and non-targeting control siRNA(#D-001210-01) were purchased from Dharmacon, Inc (Lafayette, CO). Each siRNA transfection into CaSki or other type of cells was carried out by LipoJet in vitro transfection reagent (SignaGen, Frederick, MD, USA). Plasmid pHBL03 has a short isoform of lnc-FANCI-2 cDNA (lnc-FANCI-2a-PA1, GenBank Acc. No. MT669801). Overlapping PCR products generated from 5’ and 3’ RACE PCR products with a primer pair of oHBL 17 and oHBL 18 were digested and inserted into pcDNA3 at Hind III and Xba I sites. The insertion was verified by sequencing. Plasmid pHBL05 has a long isoform of lnc-FANCI-2 (lnc-FANCI-2a-PA2, GenBank Acc. No. MT669800). PCR products generated from genomic DNA with oHBL 19 and oHBL 23 were digested and swapped into pHBL03 to replace lnc-FANCI-2a-PA1 at Bsmb I and Xba I sites.

CRISPR/Cas9 system was used to knockout lnc-FANCI-2 gene from CaSki cells. Two lnc-FANCI-2 specific guide RNAs (gRNAs) were designed by an on-line CRISPR design tool developed by the Zhang lab (http://crispr.mit.edu). A modified cloning strategy was used to clone the two gRNA into pSpCas9(BB)-2A-Puro vector (Addgene, <u>Watertown, MA</u>, #62988) and described in our previous publication [42]. This strategy allows all three components (two gRNAs and Cas9) express from the same plasmid to ensure the transfected cells expressing two gRNAs simultaneously knocking out the targeted gene [42]. The sequences of primers to make guide RNAs were listed in Supplementary Table S5.

### Knockout of lnc-FANCI-2 gene in CaSki cells, PCR screening and genotyping

Two gRNAs each under control by a U6 promoter in a pSpCas9(BB)-2A-Puro vector were used to create deletion in CaSki cells. Plasmid pHBL25 which contains gRNA 1 and 6 was used to delete the entire promoter region of lnc-FANCI-2 and plasmid pHBL27 which contains gRNA 3 and 4 was used to delete only two YY1-binding motifs spanning over an 87-bp region in the lnc-FANCI-2 promoter. CaSki cells transfected with the gRNA-expressing vectors were then under 2 weeks of puromycin selection (0.1 µg/ml). The puromycin-resistant CaSki cells were diluted to a final concentration of 0.5 cells per 100 μl, plated 100 μl of diluted cells into each well of a 96-well plate, and grew under continuous purymycin selection. The colonies were inspected to ensure the clones were grown from one cell, and refed or re-plated as needed in 3 weeks of expansion.

A direct PCR was performed for number of single cell clones as described [42]. Briefly, several hundreds of cells expanded from a single cell clone were resuspended in 50 μl of phosphate-buffered saline (PBS) and then frozen-thawed on dry ice for three times. Cell lysates were treated with 4 μl of Qiagen protease (QIAGEN, Hilden, Germany, #1017782,) at 56°C for 10 min, and the protease was inactivated at 95°C for 5 mins. The resulting product was directly used for PCR screening. P1 + P5 were used for verification of promoter deletion and P3 + P4 for YY1 motifs deletion. To ensure the selected single cell clones displaying homozygous knockout of lnc-FANCI-2, total cell DNA was isolated using a QIAamp DNA Blood Minikit (QIAGEN, Hilden, Germany, #51106) and then the homozygous deletion was screened and the cells with homozygous KO were selected and confirmed by PCR. The primers used for genotyping are listed in supplementary Table S5. At least two single cell clones either with lnc-FANCI-2 promoter deletion or with YY1 motif deletion were selected. Three single cell clones from the cells transfected with an empty vector, pSpCas9(BB)-2A-Puro, were also selected and served as controls for the genotyping screening.

### Cell proliferation and viability assay

Cell proliferation and viability were determined by a CCK-8 assay (Dojindo Molecular Technologies, Rockville, MD). For proliferation assay, 500 µl of cell suspension (50,000 cells/well) were dispensed into individual wells in a 24-well plate. Cells were then treated with inhibitors (20 μM LY294002, or 10 μM U0126 respectively) after 24 h culture. At each time point of 24 h, 48 h and 72 h, 50 µl of CCK-8 solution was added in three wells of each group and incubated for 1 h in a cell culture incubator at 37°C. Cell proliferation was determined by absorbance at 450 nm, and normal medium was used to subtract background. Percentage of cell viability was calculated by the following formula: cell viability % = OD450 of inhibitors treated sample / OD450 of untreated sample × 100%.

### Colony formation, cell migration assay and senescence analysis

CaSki Cells were seeded at 1 × 10^4^, 5 × 10^3^ or 1 × 10^3^ cells/well in 6-well plates. After 24 h of cell growth, the culture medium in each well was replaced by DMEM containing 1% methylcellulose and 10% FBS. The plates were fixed with 3.7% formaldehyde and stained with 1% crystal violet in 2 weeks. Transwell cell migration assay was performed as previously described [93]. Briefly, CaSki cells were re-suspended in serum free DMEM containing 0.1% BSA. 1 × 10^5^ cells in 100 ul were then plated on top of the filter membrane in a transwell insert (SARSTEDT, Nümbrecht, GERMANY, # 83.3932.800) and incubate for 10 min at 37 °C to allow the cells to settle down, then 600 μl of complete media containing 10% FBS were added into the bottom of the lower chamber in a 24-well plate. After 24 h in culture, the cells on the top of the insert membrane were removed, and cells on the bottom were fixed, stained by 0.2% crystal violet and then counted.

Cell senescence was determined by Cellular Senescence Assay (MilliporeSigma, Burlington, MA, # QIA117-1KIT). Briefly, CaSki Cells were plated at 2 × 10^5^ cells/well in 6-well plates and cultured for 2 days to 50% confluence. Cells were washed once with 2 ml PBS and fixed with 0.5 ml of Fixative Solution at room temperature for 10 to 15 min. Then, cells were washed twice with 2 ml PBS, and 1 ml Staining Solution was added to each well. Cells were incubated at 37°C overnight without CO_2_ and protected from the light. Blue stained cells were then counted under light microscopy.

### Nuclear and cytoplasmic fractionation

CaSki and SiHa cells were fractionated by using Nuclei EZ Prep Kit (Sigma-Aldrich, #NUC-101) following the manufacturer’s protocol. The details of the method could be found in our previous publication [94]. Western blot analysis for nuclear protein SRSF3 and cytoplasmic protein GAPDH was used to determine the fractionation efficiency. The fractionated cytoplasmic and nuclear RNAs were used for detection of lnc-FANCI-2 RNA with human GAPDH RNA serving as a control for RNA fractionation efficiency of the cytoplasmic RNAs.

### RT-qPCR

Detection of lnc-FANCI-2 by RT-qPCR was performed as described [8]. Pre-designed primers for lnc-FANCI-2 are listed in Table S5. Briefly, 2 µg total RNA was converted to cDNA using Superscript First-stand Synthesis kit (Thermo Fisher Scientific, Waltham, MA, #11904018).

qPCR was performed using TaqMan gene expression Master Mix (Applied Biosystems, Waltham, MA, #4369016) on a StepOne Plus Real-Time PCR system (Applied Biosystems). TaqMan Gene Expression Assay (Applied Biosystems, #4331182) for GADPH (Hs02758991_g1) was served as an internal control. Data were plotted as fold change over the control group using the 2^-ΔΔCt^ method by which the data was normalized first to the values for GAPDH and then to the median value for control samples. Data are presented as a bar graph with mean ± SD for each group.

### Northern blot

To validate lnc-FANCI-2 KO in CaSki cells, Northern blot was performed using polyA^+^ mRNA isolated from 100 µg of total CaSki RNA with PolyATtract mRNA Isolation Systems (Promega, #Z5310) or 10 µg of total CaSki total RNA as described previously [8]. Briefly, RNA samples were denatured in NorthernMax Formaldehyde loading dye (Thermo Fisher Scientific, #AM8552) at 75°C for 15 min, then separated in an 1% formaldehyde-containing agarose gel and transferred onto a GeneScreen Plus hybridization transfer membrane (Perkin Elmer, Waltham, MA, #NEF987001PK). RNAs on the membrane were crosslinked by exposing to UV light, and then prehybridized with PerfectHyb Plus hybridization buffer (Sigma-Aldrich, St. Louis, MO, #H7003) for 2 h at 42°C. Specific oligos against lnc-FANCI-2 or GAPDH listed in Table S5 were labeled with [γ-^32^P]-ATP using T4 PNK (Thermo Fisher Scientific, Waltham, MA, #18004–010) and added into hybridization buffer for overnight incubation at 42°C. The membrane was then washed with 2× SSPE/0.5% SDS solution for 5 mins, followed by twice washes with 0.5× SSPE/0.5% SDS each for 15 min at 42°C and then exposed to a PhosphorImager screen.

### RNA-seq and data analysis

Total RNA was from WT CaSki cells, ΔYY1-D5 and ΔPr-A9 cells were extracted using TRIzol reagent (Thermo Fisher Scientific, Waltham, MA, #15596018). Total ribo-minus RNA-seq libraries, four samples in each group, were prepared with TruSeq Stranded Total RNA Library Kit and then subjected to pair-end sequencing using Illumina-HiSeq3000/4000 platform.

The obtained reads were processed using the CCBR Pipeliner utility (https://github.com/CCBR/Pipeliner). Briefly, reads were trimmed from adapters and low-quality bases using Cutadapt (version 1.18) (https://bioweb.pasteur.fr/packages/pack@cutadapt@1.18) before alignment to the custom reference genome described below. The transcripts were aligned using STAR v2.5.2b in two-pass mode [95]. Expression levels were quantified using RSEM (RNA-Seq by Expectation-Maximization) (version 1.3.0) [96] with a custom gene annotation described below.

The custom reference genome allowing quantification of both HPV16 and host expression used in this alignment consisted of the human reference genome (hg38/Dec. 2013/GRCh38) with a HPV16 FASTA (https://pave.niaid.nih.gov/) sequence added as an additional pseudochromosome. The custom gene annotation used for gene expression quantification consisted of a concatenation of the hg38 GENCODE annotation version 30 [97] and the HPV16 genome, with one other notable alteration. The hg38 v30 gene annotation is incorrect at the location of the lnc-FANCI-2 locus in the hg38 genome at chr15:89378104-89398487, which affects quantification of this gene. To correct this annotation and provide accurate quantification, therefore, we first performed BLAST against hg38 using the sequences of the 14 known isoform transcripts of lnc-FANCI-2 (MT669800 for lnc-FANCI-2a-PA2, MT669801 for lnc-FANCI-2a-PA1, MT669802 for lnc-FANCI-2b-PA2, MT669803 for lnc-FANCI-2b-PA1, MT669804 for lnc-FANCI-2c-PA2, MT669805 for lnc-FANCI-2c-PA1, MT669806 for lnc-FANCI-2d-PA2, MT669807 for lnc-FANCI-2d-PA1, MT669808 for lnc-FANCI-2e-PA2, MT669809 for lnc-FANCI-2e-PA1, MT669810 for lnc-FANCI-2f-PA2, MT669811 for lnc-FANCI-2f-PA1, MT669812 for lnc-FANCI-2g-PA2, and MT669813 for lnc-FANCI-2g-PA1). The results were used to determine the precise exon start and stop sites for each of the isoforms. These were then used to manually adjust the default hg38 v30 gene annotations at the locus indicated above to account for the correct structure of the lnc-FANCI-2 locus and its 14 isoforms.

Downstream analysis and visualization were performed within the NIH Integrated Analysis Portal (NIDAP) using R programs developed on the Foundry platform (Palantir Technologies). Briefly, raw counts data produced by RSEM were imported into the NIDAP platform, genes were filtered for low counts (<1 cpm) and the voom algorithm [98] from the Limma R package (version 3.40.6) [99] was used for quantile normalization and calculation of differentially expressed genes. Pre-ranked GSEA was performed using the Molecular Signatures Database version 6.2 [100, 101] and the fgsea package [102]. Raw data and the analyzed RNA-seq data supporting the findings in this study have been deposited in the NCBI GEO database (the accession#: GSE190904).

### Human Soluble Receptor Array

To detect the changes of soluble receptors expressed and released from parental WT CaSki cells and lnc-FANCI-2 KO cells, Proteome Profiler Human sReceptor (Soluble Receptor) Array (R&D Systems, Minneapolis, MN, # ARY012) was performed according to a manufacturer protocol. Briefly, 7 × 10^6^ WT CaSki cells and ΔPr-A9 cells were cultured in 15 ml of DMEM supplemented with 10% FBS in a T75 flask for 24 h. Cell culture supernatants were then collected and centrifuged at 4,000 × g for 15 min to remove cell debris. Cells were rinsed by 10 ml of PBS and solubilized in Lysis Buffer 17 at 4°C for 30 min. The cell lysis was then centrifuged at 14,000 × g for 5 min to remove cell debris. Protein concentration of cell lysis was measured by Micro BCA Protein Assay (PIERCE, Waltham, MA, #23235). Next, two sets of N and C membranes were blocking by Buffer 8/1 in 4-Well Multi-dish for 1 h on a rocking platform. 500 μl of culture supernatant or 100 μg cell lysate generated from the WT CaSki cells or ΔPr-A9 cells were diluted in 3 ml of Buffer 8/1 and then applied to each membrane after three washes with 1× Wash Buffer. The membranes were incubated overnight at 4 °C on a rocking platform and washed three times to remove unbound materials followed by incubation with their specific cocktail of biotinylated detection antibodies for 2 h at room temperature on a rocking platform. 2 ml of diluted Streptavidin-HRP was then applied to each membrane for 30 min at room temperature on a rocking platform after three times of wash. 1 ml of the prepared Chemi Reagent Mix was added onto the membranes after three times of wash. The signal was then captured by ChemiDoc Touch Imaging System (Bio-Rad). Quantification of the relative protein expression in ΔPr-A9 cells compared to the WT cells was determined in Image Lab software (Bio-Rad). The average signal of duplicate spots subtracting background value from negative control spots represented the levels of each protein. The relative change in protein levels was then determined by comparing ΔPr-A9 cells to the WT cells.

### Antibodies and Immunoblot

Immunoblot analysis was performed for individual proteins by using the following antibodies: anti-MCAM (#17564-1-AP), anti-E2F1 (#12171-1-AP), anti-MAP4K4 (#55247-1-AP), and anti-p53 (#10442-1-AP) antibodies were from ProteinTech (Rosemont, IL). Anti-tubulin (#T5201) antibodies were from Sigma-Aldrich (St. Louis, MO). Anti-hnRNP C1/C2 (#Ab10294) and anti-RAC3 (#ab129062) antibodies were from Abcam (Cambridge, United Kingdom). Anti-CCND2 (Cyclin D2, #3741), anti-Akt (#9272), anti-phospho-Akt (#9271), anti-Erk1/2 (p44/42 MAPK, #4695), anti-phospho-Erk1/2 (#4370), and anti-IGFBP3 (D1U9C, #25864) were from Cell Signaling Technology (Danvers, MA). Anti-VIM (#MA5-11883) was from Thermo Fisher Scientific (Waltham, MA). Anti-PODXL2 (#AF1524), anti-ECM1 (#MAB39371), anti-TIMP2 (#AF971) and anti-ADAM8 (#AF1031) were from R&D Systems (Minneapolis, MN). Anti-SRSF3 (#NBP2-76892) was from Novus Biologicals (Centennial, CO). Rabbit polyclonal anti-HPV16 E7 (#GTX133411) was from GeneTex (Irvine, CA).

### RAS GTPase Chemi ELISA

RAS GTPase activity in cell lysate of the WT CaSki or ΔPr-A9 cells was determined by RAS GTPase Chemi ELISA Kit (Active Motif, Carlsbad, CA, # 52097) according to the manufacturer protocol. Briefly, 2 × 10^7^ cells were solubilized in 500 µl of Complete Lysis/Binding buffer at 4°C for 15 min. Protein concentration of cell extract was then measured by Micro BCA Protein Assay (PIERCE, #23235) after cell debris was removed by centrifuged at 14,000 × g for 10 min. 2 µg of GST-Raf-RBD in 50 µl of Complete Lysis/Binding buffer was coated in each well of a 96-well plate by incubating for 1 h at 4°C with 100 rpm agitation. After three times of wash, 50 µg of extract in 50 µl Complete Lysis/Binding buffer was then added in each well. HeLa (EGF treated) extract was served as a positive control. The plate was incubated for 1 h at room temperature with 100 rpm agitation. The active RAS protein (GST-Raf-RBD binding) in each well was then detected by incubating with primary RAS antibody and three times of wash, and then, HRP-conjugated secondary antibody and three times of wash. Chemiluminescence in each well was read in luminometer by adding 50 µl room-temperature Chemiluminescent Working Solution.

### RNA *in situ* Hybridization (RNA-ISH) and Immunofluorescence (IF) Staining

Endogenous lnc-FANCI-2 in cervival cancer tissues and their derived cell lines was examined by RNAScope Multiplex Fluorescent V2 Assay (Advanced Cell Diagnostics, Minneapolis, MN, #323100) as described previously [8]. A custom-designed probe targeting to nt 359-1713 of GenBank Acc. No. MT669800.1 transcript for lnc-FANCI-2a-PA2 was utilized. Dual staining of lnc-FANCI-2 and IGFBP RNA was performed according to RNAscope Multplex Fluorescent v2 Manual part 2 (#323100), and the channel one probe for IGFBP3 (#310351) and channel three probe for lnc-FANCI-2 (#509061-C3) were applied.

For RNA-ISH combined IF staining of MCAM protein, 3 × 10^5^ of CaSki cells were grown on a glass coverslip in a 6-well plate for 24 h before transfection. Plasmid pHBL05 containing a long isoform of lnc-FANCI-2a-PA2 (GenBank Acc. No. MT669800.1) were transfected into cells with LipoD 293 DNA in vitro transfection reagent. Cultured Adherent Cell Sample Preparation for the RNAscope Multplex Fluorescent v2 was performed according to a manufacturer protocol (Advanced Cell Diagnostics, Minneapolis, MN, MK-50-010). Briefly, the cells were washed with PBS and fixed by 10% neutral buffered formalin at room temperature for 30 min. The cells were washed with PBS three times and dehydrated by 50% ethanol for 5 min, 70% ethanol for 5 min, 100% ethanol for 5 min, and 100% ethanol for 10 min. The cells were then rehydrated by 70% ethanol for 2 min, 50% ethanol for 2 min, and PBS for 10 min. The cells were then applied to hydrogen peroxide at RT for 10 min and washed with distilled water for three times. The cells were then applied to Protease III (1:15 dilution with PBS) at RT for 10 min. The cells were ready for RNAscope Multplex Fluorescent v2 Manual part 2 (#323100). IF was performed after RNA-ISH but before counterstaining with DAPI. The cells were washed with PBS, and then were blocked with 2% bovine serum albumin (BSA) in PBS for 1 h at 37°C or overnight at 4°C. The cells were incubated with the MCAM primary antibody (ProteinTech, Rosemont, IL, #17564-1-AP, diluted 1:200 in 2% BSA blocking buffer) for 2 h at 37°C. An Alexa Fluor-conjugated secondary antibody (1: 500, ThermoFisher Scientific) was diluted in a blocking solution and incubated for 1 h at 37°C. The slides were washed with PBS for three times. DAPI was used for nuclei counterstaining before mounting in a Prolong Gold Antifade mounting medium (Thermo Fisher Scientific, #P36934).

Confocal images were collected using a Zeiss LSM710 laser-scanning microscope equipped with a 63 × Plan-Apochromat (N.A. 1.4) objective lens. Three dimensional distributions of lnc-FANCI-2 were generated by Z stacks using ZEN 2.3 software (Zeiss).

### <u>I</u>solation of lnc-FANCI-2 <u>R</u>NA-<u>p</u>rotein <u>c</u>omplex by <u>R</u>NA <u>P</u>urification (IRPCRP) and LC-MS/MS analysis

To identify the lnc-FANCI-2 RNA-associated proteins, we pulled down lnc-FANCI-2 RNA from WT CaSki cells using IRPCRP, a modified ChIRP protocol from the published method [103], followed by mass spectrometry [60]. Briefly, a total of 30 lnc-FANCI-2 anti-sense oligo probes (oLLY496-oLLY525), each with biotinTEG at 3’ end, across entire lnc-FANCI-2 RNA (GenBank Acc. No MT669800.1) were designed using the online probe designer (singlemoleculefish.com). Two probe pools were used in IRPCRP assays: the probe pool 1 was the mixed oligos in even numbers (oLLY496 /498/500/502/504/506/508/510/512/514/516/518/520/522/524) and the probe pool 2 mixed with the oligos in odd numbers (oLLY497/499/501/503/505/507/509/511/513/515/517/519/521/523/525) (Table S5). CaSki cells were seeded and harvested at 24 h and the cell lysates from 200 million cells were used for each IRPCRP reaction. Instead of formaldehyde crosslink in the standard protocol [103], UV cross-linking (254 nm, energy at 480 mJ/cm^2^) was applied in our study to minimize non-specific binding. After UV cross-linking, the cells were lysed on ice in 1× radioimmunoprecipitation assay (RIPA) buffer (Boston Bioproducts, Ashland, MA, #BP-115) (50 mM Tris base/Tris-HCl [pH 7.4], 150 mM NaCl, 0.5% sodium deoxycholate, 0.1% sodium dodecyl sulfate [SDS], 1% NP-40) supplemented with protease inhibitors (complete mini-EDTA-free protease inhibitor cocktail, Millipore Sigma, Burlington, MA, #469315900) and RNase inhibitor (Thermo Fisher Scientific, #AM2694) for 30 min, followed by a brief sonication. The obtained cell lysates were spin for 10 min of centrifugation at 20,000 × g at 4°C and the collected cell lysate supernatants were pre-absorbed with C-1 magnetic beads (Thermo Fisher Scientific, #65002) for 1 h at room temperature before proceeding to oligo probe hybridization.

Two different hybridization protocols were used to pull down lnc-FANCI-2 RNA. One had the pre-absorbed cell lysate incubated with two separate probe pools in hybridization buffer (750 mM NaCl, 0.1% SDS, 50 mM Tris-Cl pH 7.0, 1 mM EDTA, 15% formamide) at room temperature overnight and the pre-washed C-1 magnetic beads was then added and incubated for additional 4 h; the other had two separate oligo probe pools immobilized onto pre-washed C-1 magnetic beads first at room temperature for 1 h and then mixed with the pre-absorbed cell lysates in hybridization buffer overnight at room temperature, along with the negative control-beads only IRPCRP reaction without oligo probes serving as a control in the modified protocol. After hybridization, the beads were washed with 1× wash buffer (2× NaCl and sodium citrate, 0.1% SDS) for 5 times and resuspend in 1 ml lysis buffer. 100 µl beads from each IRPCRP were set aside for RNA isolation of lnc-FANCI-2 pulldown efficiency and the leftover beads were sent for mass spectrometry analysis. The input and IRPCRP RNA were isolated using a MIRNeasy mini kit (Qiagen, #217004) following the standard protocol after proteinase K (Millipore Sigma, #71049) digestion.

For mass spectrometry analysis, the IRPCRP beads were processed for trypsin/LysC digestion before submitted to Thermo Scientific Orbitrap Exploris 240 Mass Spectrometer and a Thermo Dionex UltiMate 3000 RSLCnano System for Proteinomics. Peptides from trypsin digestion were loaded onto a peptide trap cartridge at a flow rate of 5 µl/min. The trapped peptides were eluted onto a reversed-phase Easy-Spray Column PepMap RSLC, C18, 2 µM, 100A, 75 µm × 250 mm (Thermo Scientific) using a linear gradient of acetonitrile (3-36%) in 0.1% formic acid. The elution duration was 110 min at a flow rate of 0.3 µl/min. Eluted peptides from the Easy-Spray column were ionized and sprayed into the mass spectrometer, using a Nano Easy-Spray Ion Source (Thermo Fisher Scientific) under the following settings: spray voltage, 1.6 kV, Capillary temperature, 275 °C. Other settings were empirically determined. Raw data files were searched against human protein sequences database using the Proteome Discoverer 2.4 software (Thermo Fisher Scientific, San Jose, CA) based on the SEQUEST algorithm. All the protein peptides identified in the IRPCRP pulldowns were summarized in Table S4.

## ACKNOWLEDGEMENTS

We thank Louise T. Chow of the University of Alabama at Birmingham for her critical reading of the manuscript. We thank Craig Meyers of Penn State University Hershey Medical Center for providing HPV16- and HPV18-infected raft cultures, Xing Xie and Yang Li of Zhejiang University Women’s Hospital of China for cervical tissue sections, and Johannes G. Schweizer from Arbor Vita corporation for anti-HPV16 E6-specific antibodies. This study was supported by the Intramural Research Program of the NIH, National Cancer Institute, Center for Cancer Research.

## DATA AVAILABILITY

NCBI GEO database Acc. No. GSE190904 for CaSki cell RNA-seq.

## AUTHORS CONTRIBUTIONS

H. L., L. Y., and Z.M.Z. designed the study. H. L. and L. Y. performed the experiments. H. L., L. Y., V.M., T. M., M. Y., P. J., M.C., D. R. L., and Z.M.Z. analyzed data and participated in the discussion and interpretation of the results. H. L., L. Y., and Z.M.Z drafted the manuscript. H. L., L. Y., and Z.M.Z. revised the manuscript. All authors read and approved of the final manuscript.

## Supporting Tables

**Table S1.** Genome-wide Effect of lnc-FANCI-2 KO in ΔPr-A9 and ΔYY1 cells on gene expression when compared with parental WT CaSki cells.

**Table S2**. The genes with altered expression found in both ΔPr-A9 and ΔYY1-D5 cells when compared with parental WT CaSki cells.

**Table S3.** The genes with increased expression in KRAS signaling and EMT and with decreased expression in IFN-gamma and IFN-alpha responses in lnc-FANCI-2 KO cells (ΔPr-A9 and ΔYY1-D5) over the parental WT CaSki cells.

**Table S4.** CaSki proteins and their PSMs identified by lnc-FANCI-2 IRPCRP-LC-MS/MS

**Table S5**. Sequences of oligonucleotides used in this study.

## Supporting Figures

**Fig S1.**
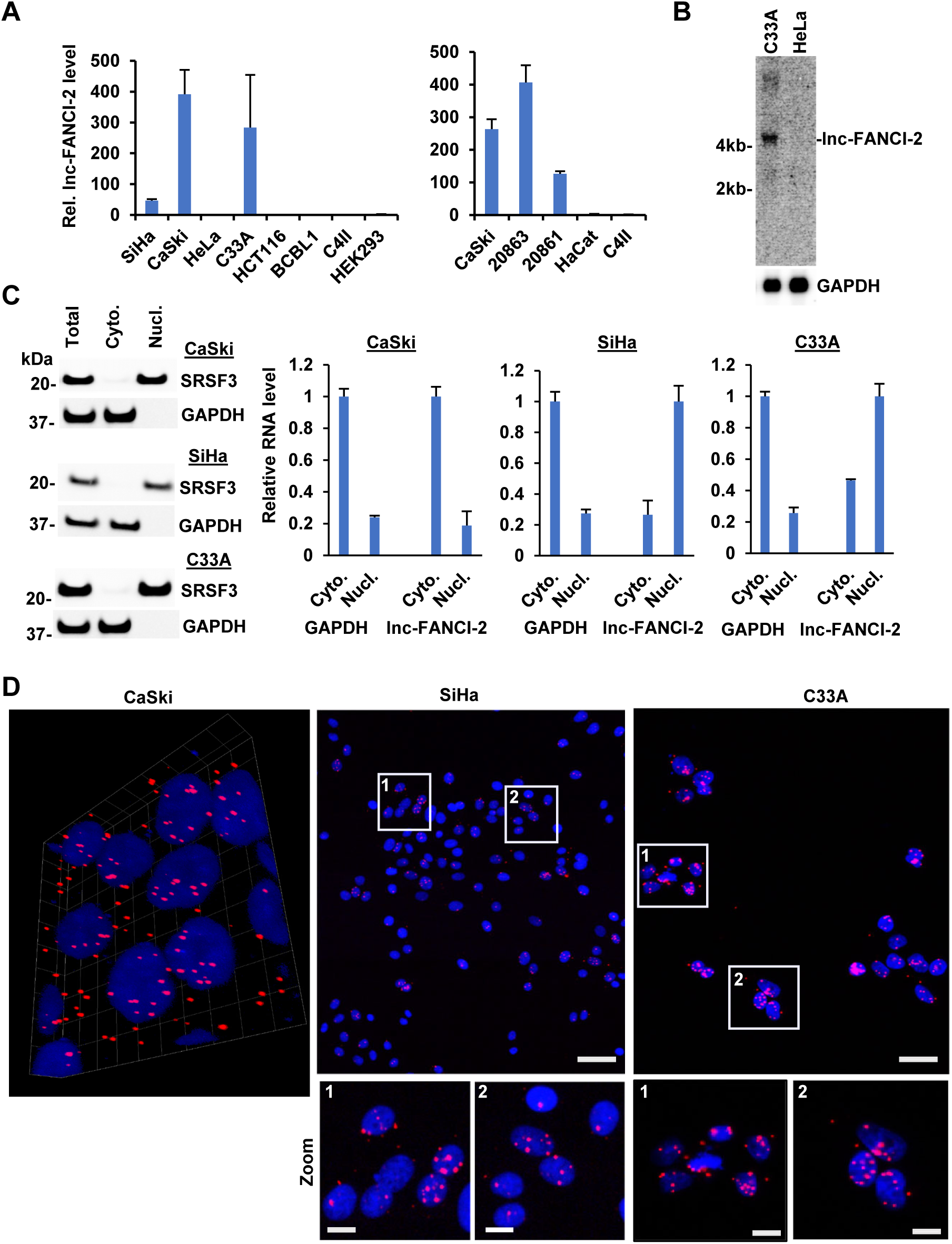
Selective expression of lnc-FANCI-2 RNA in cervical cancer cell lines. (A)) RT-qPCR detection of lnc-FANCI-2 in HPV16^+^ cervical cancer cell lines SiHa, CaSki, and W12 20861 (integrated HPV16) and 20863 (episomal HPV16), HPV18^+^ cervical cancer cell lines HeLa and C4II, and HPV^-^ cell lines C33A (cervical cancer cells with mutations of p53 and pRb), HCT116 (colorectal cancer cells), BCBL-1 (body cavity B lymphoma cells), HEK293 (Ad5 E1/E2-immortalized human kidney cells), and HaCaT (spontaneously immortalized human epidermal cells) in triplicates. (B) HeLa cells express no lnc-FANCI-2 when compared with C33A cells by Northern blot. (C) lnc-FANCI-2 is mainly cytoplasmic in CaSki but nuclear in SiHa and C33A cells. Cytoplasmic and nuclear fractionation efficiency was blotted for nuclear SRSF3 (serine- and arginine-rich splicing factor 3) and cytoplasmic GAPDH. Total fractionated cytoplasmic and nuclear RNAs were quantified for lnc-FANCI-2 by RT-qPCR in triplicates, with GAPDH RNA serving as an internal control for RNA fractionation efficiency. (D) Subcellular lnc-FANCI-2 (red) localization in CaSki, SiHa, and C33A cells determined by RNAscope RNA ISH analysis. Nuclei were stained with DAPI (blue). Scale bars: 25 μm in the top and 10 μm in the zoom.

**Fig. S2.**
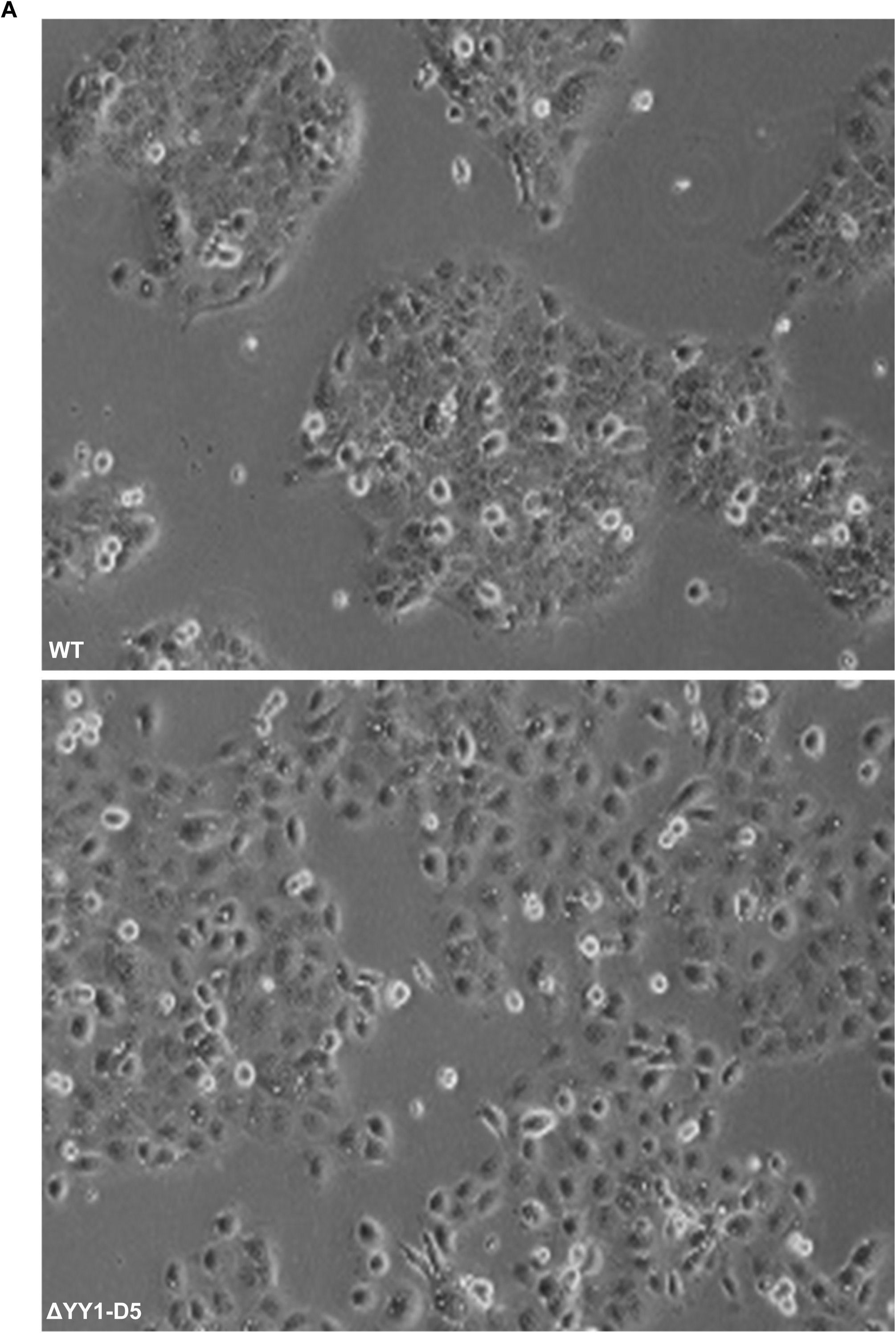
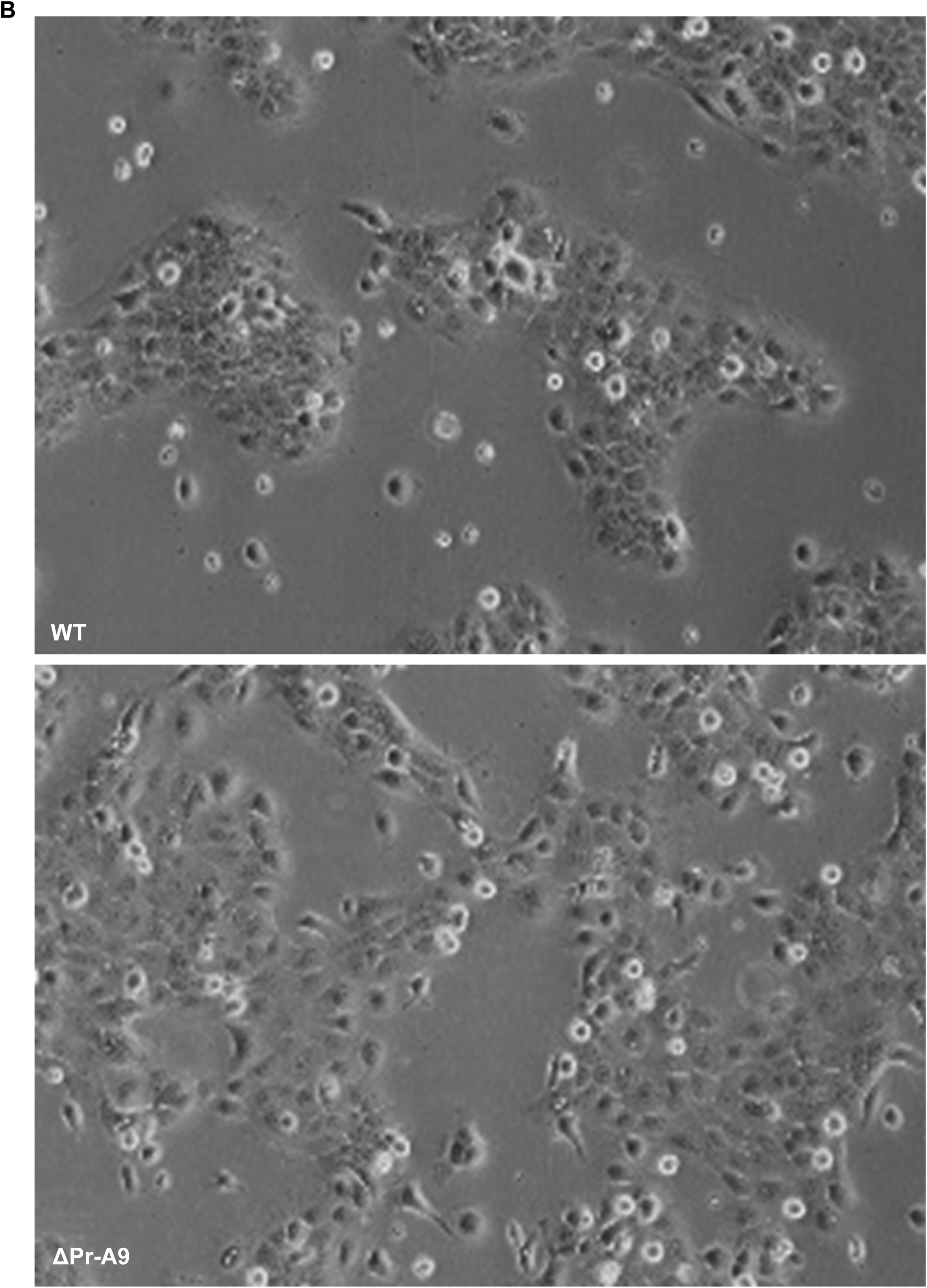
lnc-FANCI-2 KO on characteristic CaSki cell growth behavior and morphology imaged at 24 h after spreading. (A) Parental WT CaSki and ΔYY1-D5 CaSki cells. (B) Parental WT CaSki and ΔPr-A9 CaSki cells.

**Fig. S3.**
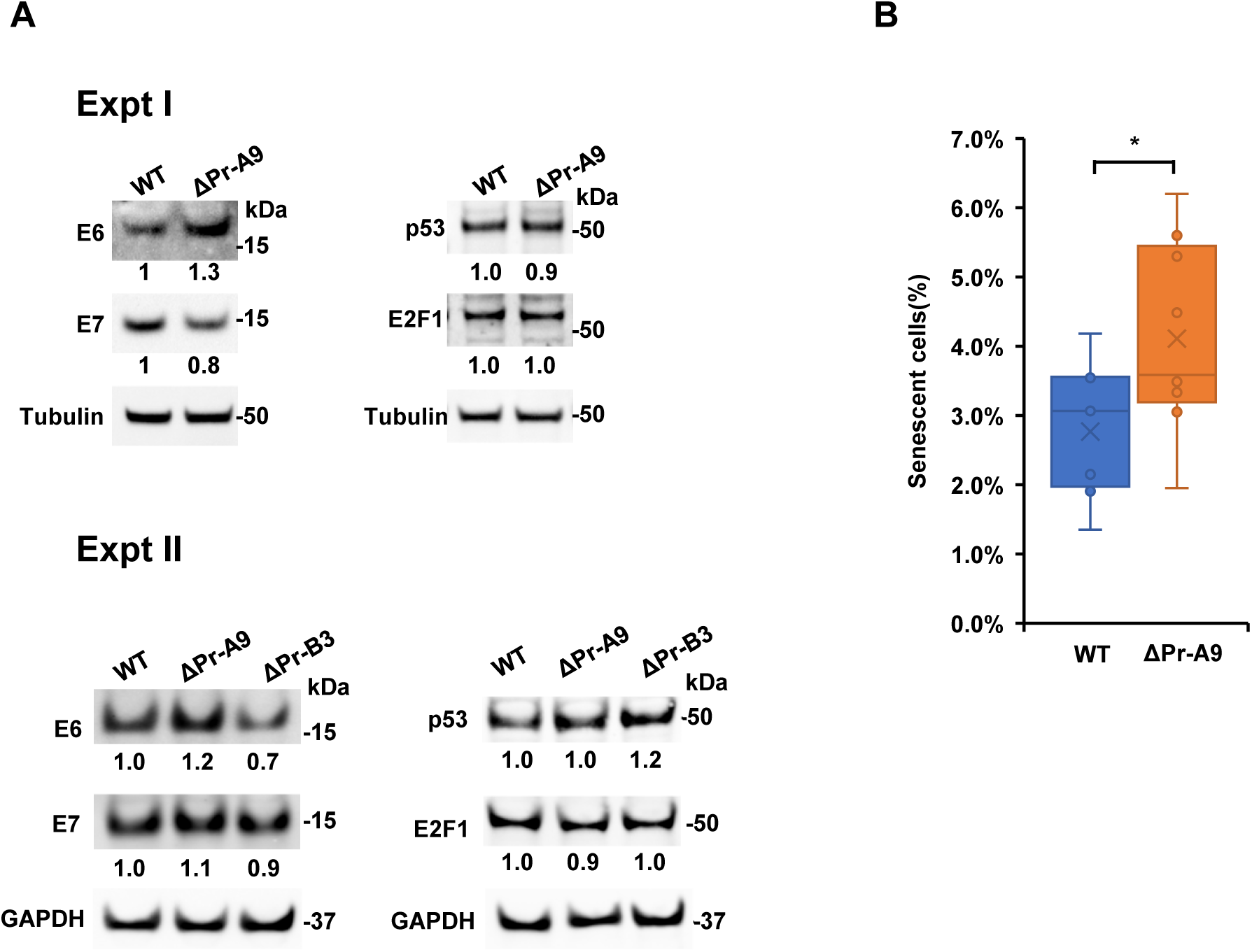
lnc-FANCI-2 KO on the expression of viral oncoproteins and cell senescence. (A) Effect of lnc-FANCI-2 KO in CaSki cells on the expression of HPV16 E6 and E7 and their downstream targets. Total cell extracts from parental CaSki, ΔPr-A9, and ΔPr-B3 cells were immunobloted with the corresponding antibodies. Tubulin or GAPDH served as a protein loading control. The relative protein levels of E6, E7, p53, and E2F1 were calculated after normalizing to tubulin or GAPDH in the corresponding experiment. Expt I or II = experiment I or II. (B) KO of lnc-FANCI-2 expression enhances cell senescence in β-gal analysis.

**Fig S4.**
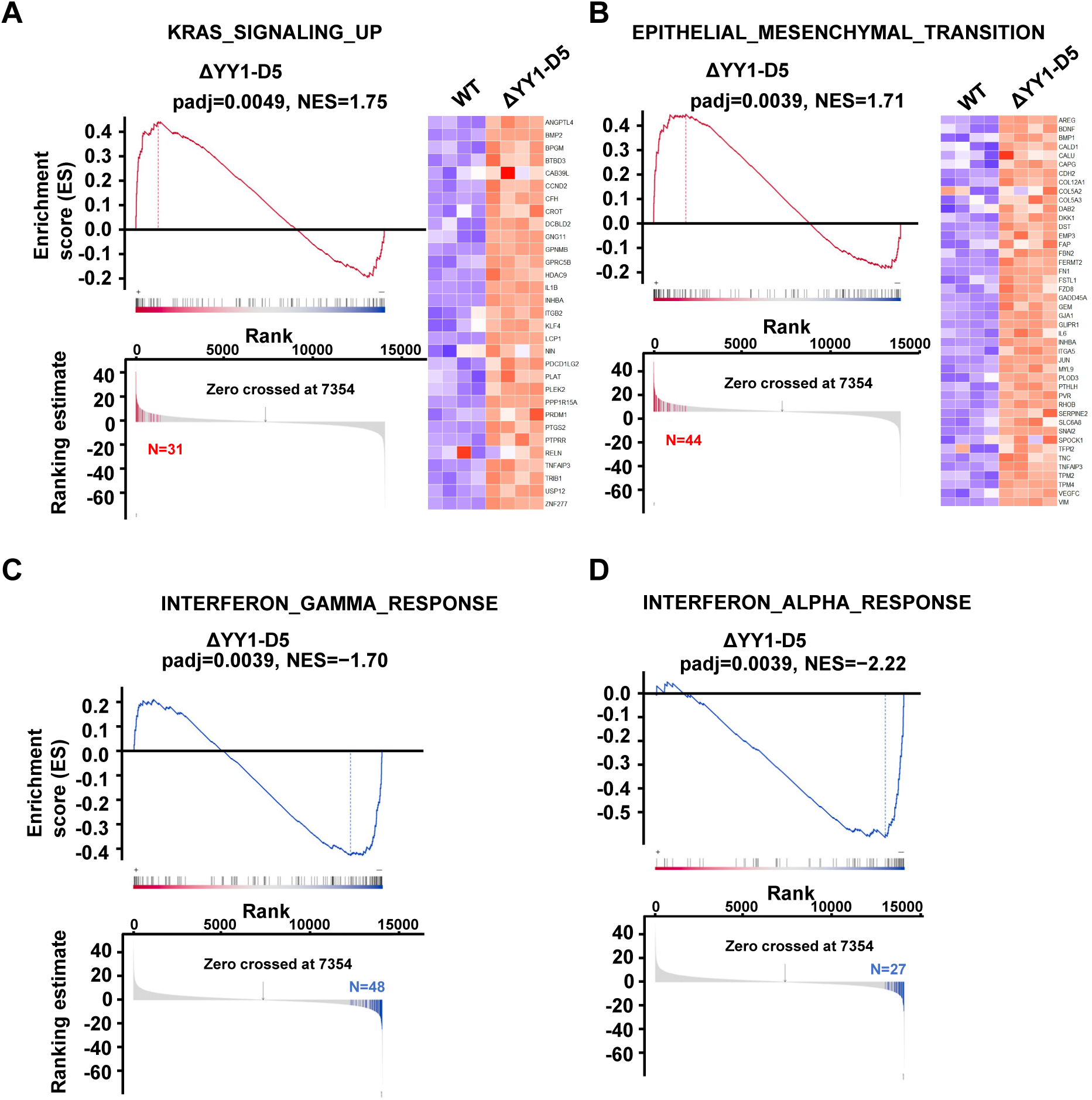
GSEA Enrichment plots show enrichment scores and gene hits enriched in ΔYY1-D5 cells using Hallmark gene sets. (A) KRAS_SIGNALING_UP. (B) EPITHELIAL_MESENCHYMAL_TRANSITION. (C) INTERFERON_GAMMA_RESPONSE. (D) INTERFERON_ALPHA_RESPONSE. The heatmaps on the right side of Enrichment plot in the panels A and B visualize the genes with differentiated expression enriched in each pathway in ΔYY1-D5 cells.

**Fig. S5.**
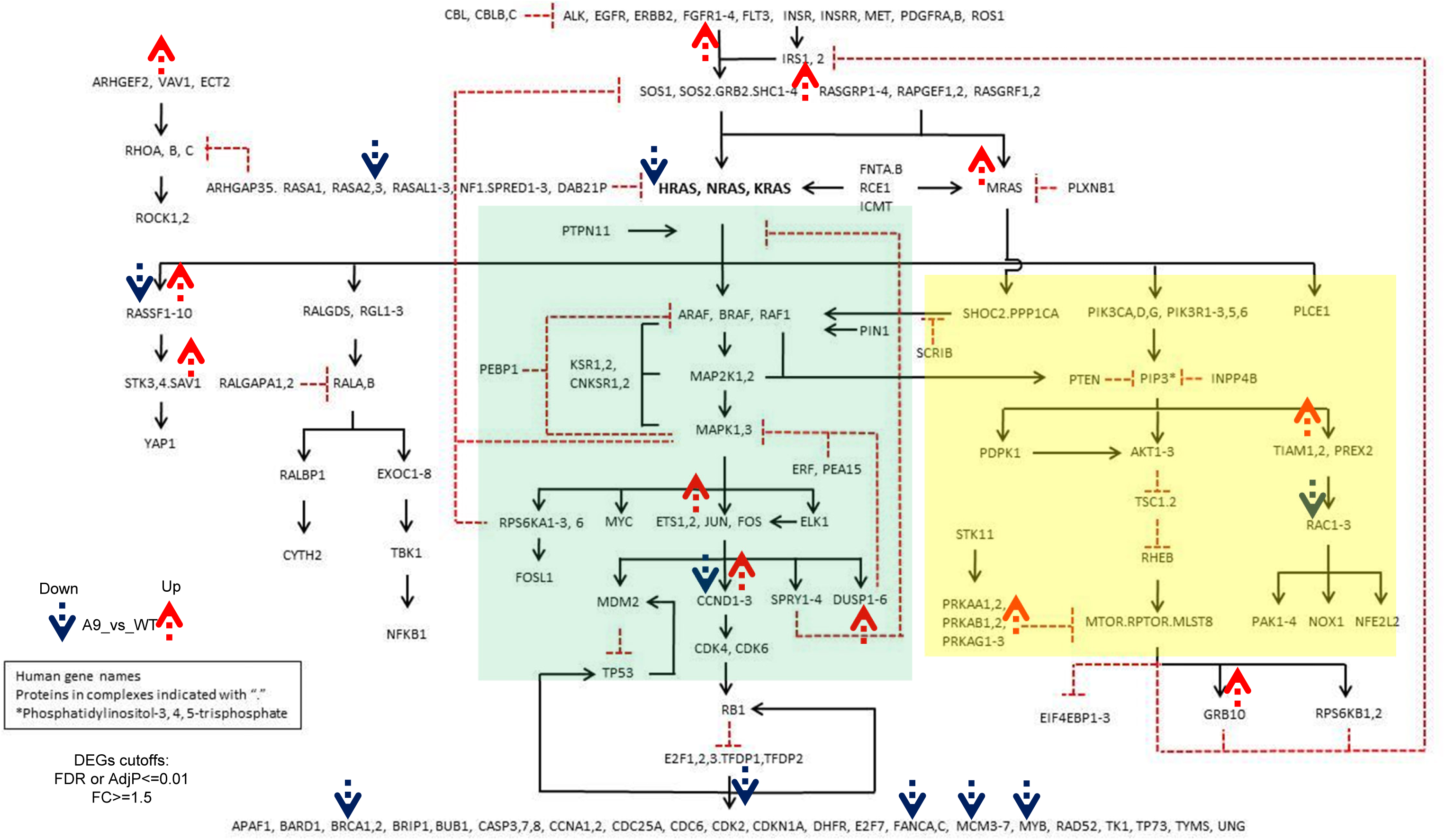
Pathway map of RAS Initiative with highlighted differentially expressed genes (DEGs) in ΔPr-A9 cells vs parental wild type cells. DEGs are mainly distributed and clustered in two main branches of RAS pathway: MAPK signaling branch and PI3K/AKT branches. DEGs were derived from RNA-seq data at cutoff of adjusted p-value <=0.01 and fold change>1.5 for both up- and down-regulated genes (indicated by red and blue arrows respectively) between ΔPr-A9 vs WT CaSki cells using three common analysis methods (DESeq2, edgeR, and limma-voom) and in-house BRB analysis pipeline from NCI. Highlighted genes in the pathway map are derived as DEGs in at least three of the analysis methods, although in most cases, these DEGs behaved consistently across all 4 methods. MAPK signaling branch is highlighted in transparent green box and PI3K/AKT branch is highlighted in transparent yellow box within the pathway map of RAS signaling pathway, which was collectively collated from community inputs that were organized and stimulated by RAS Initiative, and maintained as a common knowledge basis at URL below: https://www.cancer.gov/research/key-initiatives/ras/ras-central/blog/2015/ras-pathway-v2, which was also described in recent review (Figure 1 in Nissley and McCormick, 2022). Reference (where the Ras pathway map has been described): Nissley, D. V. and McCormick, F. (2022). RAS at 40: update from the RAS Initiative. *Cancer Discov.* https://doi.org/10.1158/2159-8290.CD-21-1554

**Fig S6.**
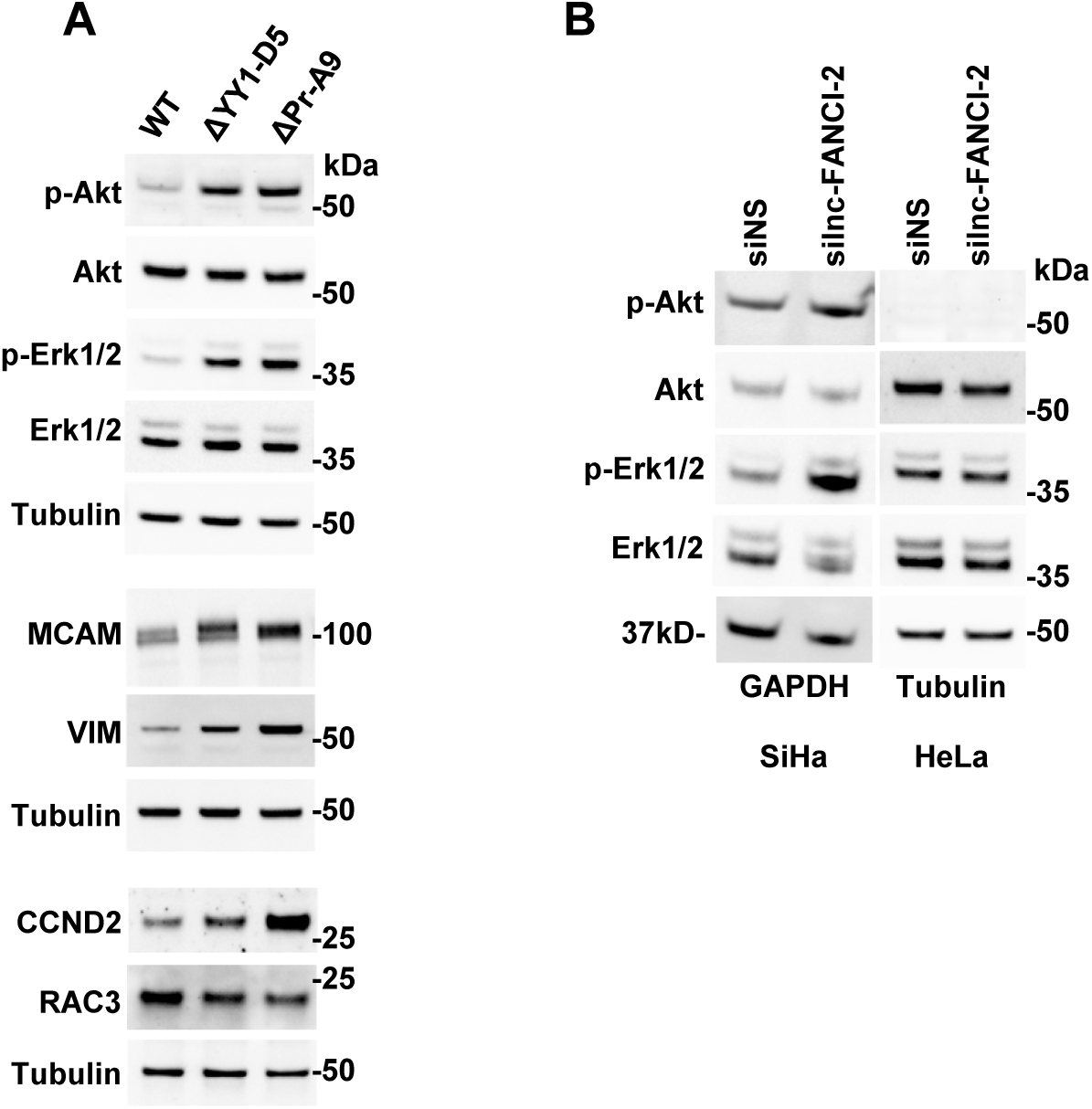
Knockout or Knockdown of lnc-FANCI-2 affects RAS signaling. (A) Selective validation of increased expression of RAS signaling-related downstream genes in ΔPr-A9 and ΔYY1-D5 cells. (B) KD of lnc-FANCI-2 expression in SiHa cells enhanced phosphorylation of Akt and Erk1/2 but did not in HeLa cells expressing no lnc-FANCI-2.

**Fig. S7.**
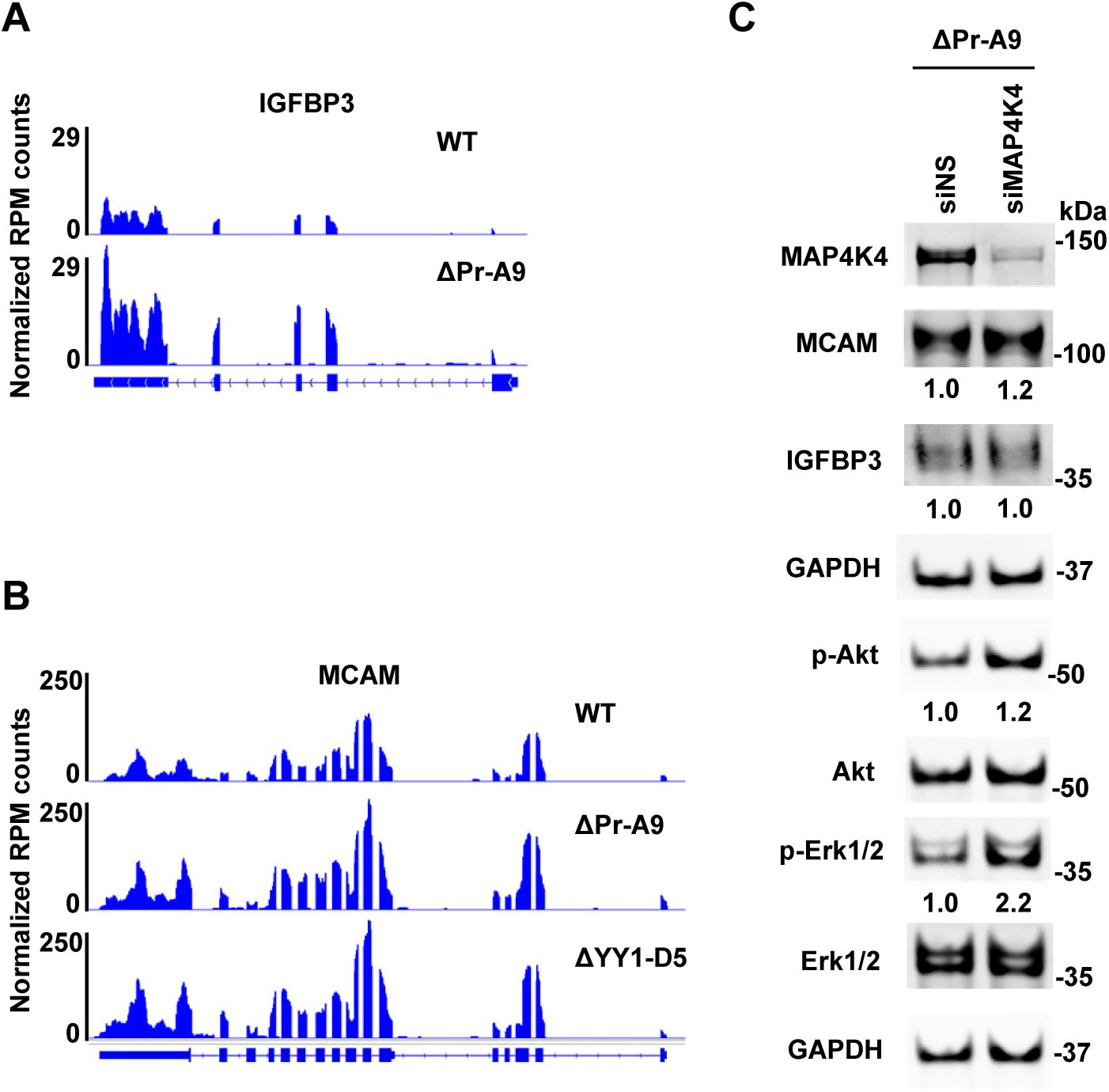
Expression of IGFBP3, MCAM, and MAP4K4 and lnc-FANCI-2. (A-B) RNA-seq reads-coverage of IGFBP3 and MCAM by IGV illustrates the increased expression of IGFBP3 and MCAM in lnc-FANCI-2 KO cells ΔPr-A9 and/or ΔYY1-D5 when compared to the parental WT CaSki cells. (C) KD of MAP4K4 expression in lnc-FANCI-2 KO ΔPr-A9 cells further enhances phosphorylation of Erk1/2 but little on MCAM and p-Akt and no effect on IGFBP3.

**Fig. S8.**
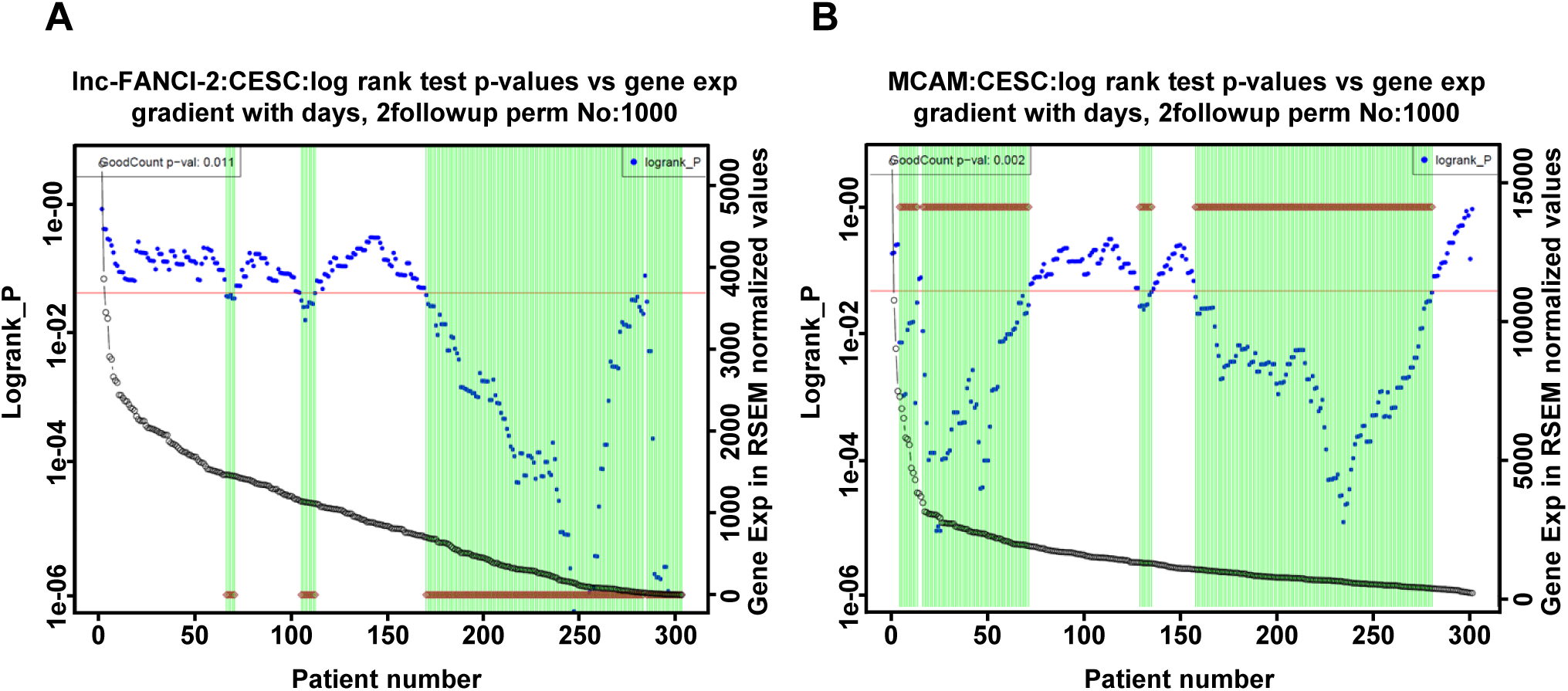
Expression levels of lnc-FANCI-2 and MCAM are significantly associated with TCGA CESC (cervical squamous cell carcinoma) cancer patients’ survival outcome. The expression of Inc-FANCI-2 (A) and of MCAM (B) along with survival of the same group of 304 cervical cancer patients were analyzed by our in-house GradientScanSurv pipeline (Yi et al 2018, PLoS One 13:e0207590). The **open grey circles** in each plot are ranked expression levels from high (left) to low (right) of Inc-FANCI-2 (A) and MCAM (B) from all patient samples as indicated by y-axis at the right side of the plot (in RSEM normalized value of RNAseq data). At each breaking point denoted by patient number (x-axis), the log rank test (y-axis at the left side of the plot) was performed between the higher expression group at the left side of the cut-point vs the lower expression group at the right side of the cut-point and the indicated log rank p-values (**blue dots**) are shown as at each cut-point at x-axis. The **horizontal red line** is the p=0.05 cutoff line for the log rank test p-values. If a log rank p-value (**blue dot**) at a cut-point is below the cutoff (p<0.05), there will be a **vertical green line** shown at the corresponding cut-point, indicative of a significant log rank test p-value, and also a **brown** diamond plotted either at the bottom part of the **vertical green line** (A) if higher expression of lnc-FANCI-2 led to less severe outcome (dying slower than the lower expressors), or at the top part of the **vertical green line** (B) if higher expression of MCAM led to more severe outcome (dying faster than the lower expressors). The original GoodCount was defined as the number of significant log rank tests (the default significance level was set as 0.05) across all possible cut-points for the original dataset. We employed the bootstrap approach to permutate the data for 1000 times in this case. A similar procedure each time was performed like the original dataset and a permutated GoodCount was obtained for each permutated dataset. The GoodCount p-value was derived as the proportion of the times that the GoodCounts of permutated datasets were no less than that of the original dataset (Yi et al 2018, PLoS One 13:e0207590) and shown at the left top corner of each plot, 0.011 for Inc-FANCI-2 in panel A and 0.002 for MCAM in panel B. A significant GoodCount p-value (**blue dots** in both panels A and B) is indicative of significant association of the expression levels with the corresponding survival outcomes of the analyzed patient samples. Not shown in the panels A and B are the coxph p-value from univariant expression-based cox regression models (coxph p-value: derived by original gene expression values; or coxph p-value by rank: derived by corresponding ranks of gene expression values) and the cut-point-specific FDR and FDR2. FDR for each cut-point was defined as the portion of cases of permutated datasets that have log rank test p-values no higher than that of original dataset. FDR2 for each cut-point was defined as the portion of cases of permutated datasets that have log rank test p-values no higher than default setting of significance as 0.05.

## REFERENCES

1. Bray F, Laversanne M, Sung H, Ferlay J, Siegel RL, Soerjomataram I, Jemal A: Global cancer statistics 2022: GLOBOCAN estimates of incidence and mortality worldwide for 36 cancers in 185 countries. CA Cancer J Clin 2024, 74:229–263.

2. Bosch FX, Lorincz A, Munoz N, Meijer CJ, Shah KV: The causal relation between human papillomavirus and cervical cancer. J Clin Pathol 2002, 55:244–265.

3. Munoz N, Bosch FX, de Sanjose S, Herrero R, Castellsague X, Shah KV, Snijders PJ, Meijer CJ, International Agency for Research on Cancer Multicenter Cervical Cancer Study G: Epidemiologic classification of human papillomavirus types associated with cervical cancer. N Engl J Med 2003, 348:518–527.

4. Walboomers JM, Jacobs MV, Manos MM, Bosch FX, Kummer JA, Shah KV, Snijders PJ, Peto J, Meijer CJ, Munoz N: Human papillomavirus is a necessary cause of invasive cervical cancer worldwide. J Pathol 1999, 189:12–19.

5. McCredie MR, Sharples KJ, Paul C, Baranyai J, Medley G, Jones RW, Skegg DC: Natural history of cervical neoplasia and risk of invasive cancer in women with cervical intraepithelial neoplasia 3: a retrospective cohort study. Lancet Oncol 2008, 9:425–434.

6. Niraj J, Farkkila A, D’Andrea AD: The Fanconi Anemia Pathway in Cancer. Annu Rev Cancer Biol 2019, 3:457–478.

7. Hoskins EE, Morris TA, Higginbotham JM, Spardy N, Cha E, Kelly P, Williams DA, Wikenheiser-Brokamp KA, Duensing S, Wells SI: Fanconi anemia deficiency stimulates HPV-associated hyperplastic growth in organotypic epithelial raft culture. Oncogene 2009, 28:674–685.

8. Liu H, Xu J, Yang Y, Wang X, Wu E, Majerciak V, Zhang T, Steenbergen RDM, Wang HK, Banerjee NS, et al: Oncogenic HPV promotes the expression of the long noncoding RNA lnc-FANCI-2 through E7 and YY1. Proc Natl Acad Sci U S A 2021, 118:e2014195118.

9. Santegoets LA, van Baars R, Terlou A, Heijmans-Antonissen C, Swagemakers SM, van der Spek PJ, Ewing PC, van Beurden M, van der Meijden WI, Helmerhorst TJ, Blok LJ: Different DNA damage and cell cycle checkpoint control in low- and high-risk human papillomavirus infections of the vulva. Int J Cancer 2012, 130:2874–2885.

10. Hoskins EE, Morreale RJ, Werner SP, Higginbotham JM, Laimins LA, Lambert PF, Brown DR, Gillison ML, Nuovo GJ, Witte DP, et al: The fanconi anemia pathway limits human papillomavirus replication. J Virol 2012, 86:8131–8138.

11. Park JW, Pitot HC, Strati K, Spardy N, Duensing S, Grompe M, Lambert PF: Deficiencies in the Fanconi anemia DNA damage response pathway increase sensitivity to HPV-associated head and neck cancer. Cancer Res 2010, 70:9959–9968.

12. Park JW, Shin MK, Lambert PF: High incidence of female reproductive tract cancers in FA-deficient HPV16-transgenic mice correlates with E7’s induction of DNA damage response, an activity mediated by E7’s inactivation of pocket proteins. Oncogene 2014, 33:3383–3391.

13. Hietanen S, Lain S, Krausz E, Blattner C, Lane DP: Activation of p53 in cervical carcinoma cells by small molecules. Proc Natl Acad Sci U S A 2000, 97:8501–8506.

14. Tommasino M, Accardi R, Caldeira S, Dong W, Malanchi I, Smet A, Zehbe I: The role of TP53 in Cervical carcinogenesis. Hum Mutat 2003, 21:307–312.

15. Prior IA, Lewis PD, Mattos C: A comprehensive survey of Ras mutations in cancer. Cancer Res 2012, 72:2457–2467.

16. Fernández-Medarde A, Santos E: Ras in cancer and developmental diseases. Genes Cancer 2011, 2:344–358.

17. Roman A, Munger K: The papillomavirus E7 proteins. Virology 2013, 445:138–168.

18. Vande Pol SB, Klingelhutz AJ: Papillomavirus E6 oncoproteins. Virology 2013, 445:115–137.

19. Liu X, Dakic A, Zhang Y, Dai Y, Chen R, Schlegel R: HPV E6 protein interacts physically and functionally with the cellular telomerase complex. Proc Natl Acad Sci U S A 2009, 106:18780–18785.

20. Klingelhutz AJ, Foster SA, McDougall JK: Telomerase activation by the E6 gene product of human papillomavirus type 16. Nature 1996, 380:79–82.

21. Storey A, Pim D, Murray A, Osborn K, Banks L, Crawford L: Comparison of the in vitro transforming activities of human papillomavirus types. EMBO J 1988, 7:1815–1820.

22. Barbosa MS, Schlegel R: The E6 and E7 genes of HPV-18 are sufficient for inducing two-stage in vitro transformation of human keratinocytes. Oncogene 1989, 4:1529–1532.

23. Hawley-Nelson P, Vousden KH, Hubbert NL, Lowy DR, Schiller JT: HPV16 E6 and E7 proteins cooperate to immortalize human foreskin keratinocytes. EMBO J 1989, 8:3905–3910.

24. Munger K, Phelps WC, Bubb V, Howley PM, Schlegel R: The E6 and E7 genes of the human papillomavirus type 16 together are necessary and sufficient for transformation of primary human keratinocytes. J Virol 1989, 63:4417–4421.

25. Halbert CL, Demers GW, Galloway DA: The E7 gene of human papillomavirus type 16 is sufficient for immortalization of human epithelial cells. J Virol 1991, 65:473–478.

26. Phelps WC, Yee CL, Munger K, Howley PM: The human papillomavirus type 16 E7 gene encodes transactivation and transformation functions similar to those of adenovirus E1A. Cell 1988, 53:539–547.

27. DiPaolo JA, Woodworth CD, Popescu NC, Notario V, Doniger J: Induction of human cervical squamous cell carcinoma by sequential transfection with human papillomavirus 16 DNA and viral Harvey ras. Oncogene 1989, 4:395–399.

28. Schreiber K, Cannon RE, Karrison T, Beck-Engeser G, Huo D, Tennant RW, Jensen H, Kast WM, Krausz T, Meredith SC, et al: Strong synergy between mutant ras and HPV16 E6/E7 in the development of primary tumors. Oncogene 2004, 23:3972–3979.

29. Narisawa-Saito M, Yoshimatsu Y, Ohno S, Yugawa T, Egawa N, Fujita M, Hirohashi S, Kiyono T: An in vitro multistep carcinogenesis model for human cervical cancer. Cancer Res 2008, 68:5699–5705.

30. Conover CA, Bale LK, Durham SK, Powell DR: Insulin-like growth factor (IGF) binding protein-3 potentiation of IGF action is mediated through the phosphatidylinositol-3-kinase pathway and is associated with alteration in protein kinase B/AKT sensitivity. Endocrinology 2000, 141:3098–3103.

31. Allard JB, Duan C: IGF-Binding Proteins: Why Do They Exist and Why Are There So Many? Front Endocrinol (Lausanne) 2018, 9:117.

32. Bach LA: IGF-binding proteins. J Mol Endocrinol 2018, 61:T11–t28.

33. Song S, Pitot HC, Lambert PF: The human papillomavirus type 16 E6 gene alone is sufficient to induce carcinomas in transgenic animals. J Virol 1999, 73:5887–5893.

34. Herber R, Liem A, Pitot H, Lambert PF: Squamous epithelial hyperplasia and carcinoma in mice transgenic for the human papillomavirus type 16 E7 oncogene. J Virol 1996, 70:1873–1881.

35. Song S, Liem A, Miller JA, Lambert PF: Human papillomavirus types 16 E6 and E7 contribute differently to carcinogenesis. Virology 2000, 267:141–150.

36. Riley RR, Duensing S, Brake T, Munger K, Lambert PF, Arbeit JM: Dissection of human papillomavirus E6 and E7 function in transgenic mouse models of cervical carcinogenesis. Cancer Res 2003, 63:4862–4871.

37. Brake T, Lambert PF: Estrogen contributes to the onset, persistence, and malignant progression of cervical cancer in a human papillomavirus-transgenic mouse model. Proc Natl Acad Sci U S A 2005, 102:2490–2495.

38. Kato S, Endoh H, Masuhiro Y, Kitamoto T, Uchiyama S, Sasaki H, Masushige S, Gotoh Y, Nishida E, Kawashima H, et al: Activation of the estrogen receptor through phosphorylation by mitogen-activated protein kinase. Science 1995, 270:1491–1494.

39. Statello L, Guo CJ, Chen LL, Huarte M: Gene regulation by long non-coding RNAs and its biological functions. Nat Rev Mol Cell Biol 2021, 22:96–118.

40. Liu H, Zheng ZM: Linking a nuclear lncRNA to cytoplasmic lysosome integrity and cell death. Proc Natl Acad Sci U S A 2022, 119:e2123082119.

41. Scheffner M, Münger K, Byrne JC, Howley PM: The state of the p53 and retinoblastoma genes in human cervical carcinoma cell lines. Proceedings of the National Academy of Sciences 1991, 88:5523–5527.

42. BeltCappellino A, Majerciak V, Lobanov A, Lack J, Cam M, Zheng ZM: CRISPR/Cas9-Mediated Knockout and In Situ Inversion of the ORF57 Gene from All Copies of the Kaposi’s Sarcoma-Associated Herpesvirus Genome in BCBL-1 Cells. J Virol 2019, 93:e00628–00619.

43. Schlomann U, Wildeboer D, Webster A, Antropova O, Zeuschner D, Knight CG, Docherty AJ, Lambert M, Skelton L, Jockusch H, Bartsch JW: The metalloprotease disintegrin ADAM8. Processing by autocatalysis is required for proteolytic activity and cell adhesion. J Biol Chem 2002, 277:48210–48219.

44. Conrad C, Yildiz D, Cleary SJ, Margraf A, Cook L, Schlomann U, Panaretou B, Bowser JL, Karmouty-Quintana H, Li J, et al: ADAM8 signaling drives neutrophil migration and ARDS severity. JCI Insight 2022, 7:e149870.

45. Liberzon A, Birger C, Thorvaldsdottir H, Ghandi M, Mesirov JP, Tamayo P: The Molecular Signatures Database (MSigDB) hallmark gene set collection. Cell Syst 2015, 1:417–425.

46. Downward J: Targeting RAS signalling pathways in cancer therapy. Nat Rev Cancer 2003, 3:11–22.

47. Simanshu DK, Nissley DV, McCormick F: RAS Proteins and Their Regulators in Human Disease. Cell 2017, 170:17–33.

48. Castellano E, Downward J: RAS Interaction with PI3K: More Than Just Another Effector Pathway. Genes Cancer 2011, 2:261–274.

49. Asati V, Mahapatra DK, Bharti SK: PI3K/Akt/mTOR and Ras/Raf/MEK/ERK signaling pathways inhibitors as anticancer agents: Structural and pharmacological perspectives. Eur J Med Chem 2016, 109:314–341.

50. Cuesta C, Arevalo-Alameda C, Castellano E: The Importance of Being PI3K in the RAS Signaling Network. Genes (Basel) 2021, 12:1094.

51. Vanhaesebroeck B, Perry MWD, Brown JR, Andre F, Okkenhaug K: PI3K inhibitors are finally coming of age. Nat Rev Drug Discov 2021, 20:741–769.

52. Favata MF, Horiuchi KY, Manos EJ, Daulerio AJ, Stradley DA, Feeser WS, Van Dyk DE, Pitts WJ, Earl RA, Hobbs F, et al: Identification of a novel inhibitor of mitogen-activated protein kinase kinase. J Biol Chem 1998, 273:18623–18632.

53. Serrano ML, Sánchez-Gómez M, Bravo MM: Insulin-like growth factor system gene expression in cervical scrapes from women with squamous intraepithelial lesions and cervical cancer. Growth Horm IGF Res 2007, 17:492–499.

54. Wang Z, Xu Q, Zhang N, Du X, Xu G, Yan X: CD146, from a melanoma cell adhesion molecule to a signaling receptor. Signal Transduct Target Ther 2020, 5:148.

55. Kebir A, Harhouri K, Guillet B, Liu JW, Foucault-Bertaud A, Lamy E, Kaspi E, Elganfoud N, Vely F, Sabatier F, et al: CD146 short isoform increases the proangiogenic potential of endothelial progenitor cells in vitro and in vivo. Circ Res 2010, 107:66–75.

56. Stalin J, Harhouri K, Hubert L, Garrigue P, Nollet M, Essaadi A, Muller A, Foucault-Bertaud A, Bachelier R, Sabatier F, et al: Soluble CD146 boosts therapeutic effect of endothelial progenitors through proteolytic processing of short CD146 isoform. Cardiovasc Res 2016, 111:240–251.

57. Joshkon A, Heim X, Dubrou C, Bachelier R, Traboulsi W, Stalin J, Fayyad-Kazan H, Badran B, Foucault-Bertaud A, Leroyer AS, et al: Role of CD146 (MCAM) in Physiological and Pathological Angiogenesis-Contribution of New Antibodies for Therapy. Biomedicines 2020, 8:633.

58. Baxter RC: Insulin-like growth factor binding protein-3 (IGFBP-3): Novel ligands mediate unexpected functions. J Cell Commun Signal 2013, 7:179–189.

59. Varma Shrivastav S, Bhardwaj A, Pathak KA, Shrivastav A: Insulin-Like Growth Factor Binding Protein-3 (IGFBP-3): Unraveling the Role in Mediating IGF-Independent Effects Within the Cell. Front Cell Dev Biol 2020, 8:286.

60. Chu C, Chang HY: ChIRP-MS: RNA-Directed Proteomic Discovery. Methods Mol Biol 2018, 1861:37–45.

61. Gao X, Gao C, Liu G, Hu J: MAP4K4: an emerging therapeutic target in cancer. Cell Biosci 2016, 6:56.

62. Gao X, Chen G, Gao C, Zhang DH, Kuan SF, Stabile LP, Liu G, Hu J: MAP4K4 is a novel MAPK/ERK pathway regulator required for lung adenocarcinoma maintenance. Mol Oncol 2017, 11:628–639.

63. Gonzalez-Montero J, Rojas CI, Burotto M: MAP4K4 and cancer: ready for the main stage? Front Oncol 2023, 13:1162835.

64. Patterson V, Ullah F, Bryant L, Griffin JN, Sidhu A, Saliganan S, Blaile M, Saenz MS, Smith R, Ellingwood S, et al: Abrogation of MAP4K4 protein function causes congenital anomalies in humans and zebrafish. Sci Adv 2023, 9:eade0631.

65. Roth Flach RJ, Guo CA, Danai LV, Yawe JC, Gujja S, Edwards YJ, Czech MP: Endothelial Mitogen-Activated Protein Kinase Kinase Kinase Kinase 4 Is Critical for Lymphatic Vascular Development and Function. Mol Cell Biol 2016, 36:1740–1749.

66. Castello A, Hentze MW, Preiss T: Metabolic Enzymes Enjoying New Partnerships as RNA-Binding Proteins. Trends Endocrinol Metab 2015, 26:746–757.

67. Hentze MW, Castello A, Schwarzl T, Preiss T: A brave new world of RNA-binding proteins. Nat Rev Mol Cell Biol 2018, 19:327–341.

68. Crook T, Storey A, Almond N, Osborn K, Crawford L: Human papillomavirus type 16 cooperates with activated ras and fos oncogenes in the hormone-dependent transformation of primary mouse cells. Proc Natl Acad Sci U S A 1988, 85:8820–8824.

69. Matlashewski G, Schneider J, Banks L, Jones N, Murray A, Crawford L: Human papillomavirus type 16 DNA cooperates with activated ras in transforming primary cells. EMBO J 1987, 6:1741–1746.

70. Greenhalgh DA, Wang XJ, Rothnagel JA, Eckhardt JN, Quintanilla MI, Barber JL, Bundman DS, Longley MA, Schlegel R, Roop DR: Transgenic mice expressing targeted HPV-18 E6 and E7 oncogenes in the epidermis develop verrucous lesions and spontaneous, rasHa-activated papillomas. Cell Growth Differ 1994, 5:667–675.

71. Riou G, Barrois M, Sheng ZM, Duvillard P, Lhomme C: Somatic deletions and mutations of c-Ha-ras gene in human cervical cancers. Oncogene 1988, 3:329–333.

72. Eiben GL, Velders MP, Schreiber H, Cassetti MC, Pullen JK, Smith LR, Kast WM: Establishment of an HLA-A*0201 human papillomavirus type 16 tumor model to determine the efficacy of vaccination strategies in HLA-A*0201 transgenic mice. Cancer Res 2002, 62:5792–5799.

73. Golijow CD, Mouron SA, Gomez MA, Dulout FN: Differences in K-ras codon 12 mutation frequency between “high-risk” and “low-risk” HPV-infected samples. Gynecol Oncol 1999, 75:108–112.

74. Prokopakis P, Sourvinos G, Koumantaki Y, Koumantakis E, Spandidos DA: K-ras mutations and HPV infection in cervicitis and intraepithelial neoplasias of the cervix. Oncol Rep 2002, 9:129–133.

75. Strickland SW, Vande Pol S: The Human Papillomavirus 16 E7 Oncoprotein Attenuates AKT Signaling To Promote Internal Ribosome Entry Site-Dependent Translation and Expression of c-MYC. J Virol 2016, 90:5611–5621.

76. Nieto P, Ambrogio C, Esteban-Burgos L, Gómez-López G, Blasco MT, Yao Z, Marais R, Rosen N, Chiarle R, Pisano DG, et al: A Braf kinase-inactive mutant induces lung adenocarcinoma. Nature 2017, 548:239–243.

77. Yang J, Nie J, Ma X, Wei Y, Peng Y, Wei X: Targeting PI3K in cancer: mechanisms and advances in clinical trials. Mol Cancer 2019, 18:26.

78. Saliani M, Mirzaiebadizi A, Javadmanesh A, Siavoshi A, Ahmadian MR: KRAS-related long noncoding RNAs in human cancers. Cancer Gene Ther 2022, 29:418–427.

79. Martin JL, Weenink SM, Baxter RC: Insulin-like growth factor-binding protein-3 potentiates epidermal growth factor action in MCF-10A mammary epithelial cells. Involvement of p44/42 and p38 mitogen-activated protein kinases. J Biol Chem 2003, 278:2969–2976.

80. Józefiak A, Larska M, Pomorska-Mól M, Ruszkowski JJ: The IGF-1 Signaling Pathway in Viral Infections. Viruses 2021, 13:1488.

81. Jozefiak A, Pacholska-Bogalska J, Myga-Nowak M, Kedzia W, Kwasniewska A, Luczak M, Kedzia H, Gozdzicka-Jozefiak A: Serum and tissue levels of insulin-like growth factor-I in women with dysplasia and HPV-positive cervical cancer. Mol Med Rep 2008, 1:231–237.

82. Serrano ML, Romero A, Cendales R, Sánchez-Gómez M, Bravo MM: Serum levels of insulin-like growth factor-I and -II and insulin-like growth factor binding protein 3 in women with squamous intraepithelial lesions and cervical cancer. Biomedica 2006, 26:258–268.

83. Schäfer B, Marg B, Gschwind A, Ullrich A: Distinct ADAM metalloproteinases regulate G protein-coupled receptor-induced cell proliferation and survival. J Biol Chem 2004, 279:47929–47938.

84. Ohtsu H, Dempsey PJ, Eguchi S: ADAMs as mediators of EGF receptor transactivation by G protein-coupled receptors. Am J Physiol Cell Physiol 2006, 291:C1–10.

85. Dang M, Dubbin K, D’Aiello A, Hartmann M, Lodish H, Herrlich A: Epidermal growth factor (EGF) ligand release by substrate-specific a disintegrin and metalloproteases (ADAMs) involves different protein kinase C (PKC) isoenzymes depending on the stimulus. J Biol Chem 2011, 286:17704–17713.

86. Kleino I, Järviluoma A, Hepojoki J, Huovila AP, Saksela K: Preferred SH3 domain partners of ADAM metalloproteases include shared and ADAM-specific SH3 interactions. PLoS One 2015, 10:e0121301.

87. Huang YK, Cheng WC, Kuo TT, Yang JC, Wu YC, Wu HH, Lo CC, Hsieh CY, Wong SC, Lu CH, et al: Inhibition of ADAM9 promotes the selective degradation of KRAS and sensitizes pancreatic cancers to chemotherapy. Nat Cancer 2024, 5:400–419.

88. Sun C, Wu MH, Guo M, Day ML, Lee ES, Yuan SY: ADAM15 regulates endothelial permeability and neutrophil migration via Src/ERK1/2 signalling. Cardiovasc Res 2010, 87:348–355.

89. Anfosso F, Bardin N, Frances V, Vivier E, Camoin-Jau L, Sampol J, Dignat-George F: Activation of human endothelial cells via S-endo-1 antigen (CD146) stimulates the tyrosine phosphorylation of focal adhesion kinase p125(FAK). J Biol Chem 1998, 273:26852–26856.

90. Dye DE, Karlen S, Rohrbach B, Staub O, Braathen LR, Eidne KA, Coombe DR: hShroom1 links a membrane bound protein to the actin cytoskeleton. Cell Mol Life Sci 2009, 66:681–696.

91. Bardin N, Frances V, Combes V, Sampol J, Dignat-George F: CD146: biosynthesis and production of a soluble form in human cultured endothelial cells. FEBS Lett 1998, 421:12–14.

92. Xu J, Liu H, Yang Y, Wang X, Liu P, Li Y, Meyers C, Banerjee NS, Wang HK, Cam M, et al: Genome-Wide Profiling of Cervical RNA-Binding Proteins Identifies Human Papillomavirus Regulation of RNASEH2A Expression by Viral E7 and E2F1. mBio 2019, 10:e02687–18.

93. Justus CR, Leffler N, Ruiz-Echevarria M, Yang LV: In vitro cell migration and invasion assays. J Vis Exp 2014, 88:51046.

94. Yu L, Zheng ZM: Human Papillomavirus Type 16 Circular RNA Is Barely Detectable for the Claimed Biological Activity. mBio 2022, 13:e0359421.

95. Dobin A, Davis CA, Schlesinger F, Drenkow J, Zaleski C, Jha S, Batut P, Chaisson M, Gingeras TR: STAR: ultrafast universal RNA-seq aligner. Bioinformatics 2013, 29:15–21.

96. Li B, Dewey CN: RSEM: accurate transcript quantification from RNA-Seq data with or without a reference genome. BMC Bioinformatics 2011, 12:323.

97. Harrow J, Frankish A, Gonzalez JM, Tapanari E, Diekhans M, Kokocinski F, Aken BL, Barrell D, Zadissa A, Searle S, et al: GENCODE: the reference human genome annotation for The ENCODE Project. Genome Res 2012, 22:1760–1774.

98. Law CW, Chen Y, Shi W, Smyth GK: voom: Precision weights unlock linear model analysis tools for RNA-seq read counts. Genome Biol 2014, 15:R29.

99. Smyth GK: Linear models and empirical bayes methods for assessing differential expression in microarray experiments. Stat Appl Genet Mol Biol 2004, 3:Article3.

100. Liberzon A, Subramanian A, Pinchback R, Thorvaldsdottir H, Tamayo P, Mesirov JP: Molecular signatures database (MSigDB) 3.0. Bioinformatics 2011, 27:1739–1740.

101. Subramanian A, Tamayo P, Mootha VK, Mukherjee S, Ebert BL, Gillette MA, Paulovich A, Pomeroy SL, Golub TR, Lander ES, Mesirov JP: Gene set enrichment analysis: a knowledge-based approach for interpreting genome-wide expression profiles. Proc Natl Acad Sci U S A 2005, 102:15545–15550.

102. Korotkevich G, Sukhov V, Budin N, Shpak B, Artyomov MN, Sergushichev A: Fast gene set enrichment analysis. bioRxiv 2021.

103. Chu C, Quinn J, Chang HY: Chromatin isolation by RNA purification (ChIRP). J Vis Exp 2012:3912.

